# Molecular Investigations into the Unfoldase Action of Severing Enzymes on Microtubules

**DOI:** 10.1101/2019.12.11.873273

**Authors:** Rohith Anand Varikoti, Amanda C. Macke, Virginia Speck, Jennifer L. Ross, Ruxandra I. Dima

## Abstract

Microtubules associated proteins regulate the dynamic behavior of microtubules during cellular processes. Microtubule severing enzymes are the associated proteins which destabilize microtubules by removing subunits from the lattice. One model for how severing enzymes remove tubulin dimers from the microtubule lattice is by unfolding its subunits through pulling on the carboxy-terminal tails of tubulin dimers. This model stems from the fact that severing enzymes are AAA+ unfoldases. To test this mechanism, we apply pulling forces on the carboxy-terminal regions of microtubule subunits using coarse grained molecular simulations. In our simulations we used different microtubule lattices and concentrations of severing enzymes. We compare our simulation results with data from *in-vitro* severing assays and find that the experimental data is best fit by a model of cooperative removal of protofilament fragments by severing enzymes which depends on the severing enzyme concentration and placement on the microtubule lattice.

## Introduction

Microtubules (MTs) are filamentous biopolymers responsible for proper intracellular organization and cellular functions that depends on the developmental stage and function of the cell. MTs alter their activities in cells using an array of ∼ 300 accessory proteins called microtubule-associated proteins (MAPs) [Sato-Harada, et al., 1996]. Arguably one of the most important tasks undertaken by MTs is the rearrangement of the chromosomes during cell division where MTs make up the cell division machinery, called the mitotic spindle [Bailey et al., 2016]. MTs are able to push and pull during their re-organization because they are long, hollow cylinders composed of α-β-tubulin heterodimers. Tubulin dimers bind together longitudinally in a lattice of parallel protofilaments (PFs) rolled into a tube with an overall polarity where the α-tubulin is exposed at the “minus end” and the β-tubulin is exposed at the “plus end.” The filament has unique abilities to grow and shrink because it is composed of tubulin dimers, which are enzymes that change shape upon the hydrolysis of GTP.

Although the overall shape of the MT is a closed cylinder, it has long been known that the dimers can join together in a variety of lattices with various neighboring schemes, chiralities, and edge defects. For instance, PFs join laterally such that like monomers are making contacts with each other until the first PF forms a heterotypic interface with the last PF resulting in α monomers being in contact with β monomers referred to as the seam [Sui & Downing, 2010; Bailey et al., 2016; Harris, et al., 2018]. The lattice is defined by the number of PFs, isoform of tubulin, various nucleation factors, and post translational modifications, all of which are dependent on the cell type as well as the species [Sui & Downing, 2010; Chaaban & Brouhard, 2017]. The canonical 13 PF MT, is thought to be evolutionarily advantageous for efficient motor navigation and organelle transportation due to the straight geometry of the incorporated PFs [Tilney, 1973]. The divergences between different tubulin isoforms are primarily in the carboxyl-terminal tails (CTTs), an unstructured, highly charged region that sticks out from the MT surface and facilitates the interactions between many MAPs and MTs [Wehenkel & Janke, 2014; Zehr et al., 2017; Ti et al., 2018].

MAPs alter MT properties such as nucleation, growth, stabilization, and depolymerization. Enzymes that sever MTs, called MT-severing enzymes, are hexameric ATPases associated with cellular activities (AAA+) enzymes [Hartman & Vale, 1999; Vale, 2000; Frickey & Lupas, 2004; Roll-Mecak & Vale, 2008; Eckert et al., 2012]. There are three closely associated families of MT severing enzymes: katanin, spastin, and fidgetin. They share similar structures and sequence motifs in the C-terminal AAA domains used to associate to MTs, but have different N-terminal domains [Vale, 2000; Zhang, 2007]. MT-severing enzymes can only sever MTs when the CTT of tubulin is present, implying that the tail is required for severing activity [Roll-Mecak & Vale, 2008; Bailey et al., 2015; McKenney et al., 2016; Belonogov et al., 2019]. The severing action is a mechanical process where the chemical energy released by the hydrolysis of ATP is converted to mechanical work. The mechanism for how the motor interacts with the MT to sever is still highly debated.

The two theories that describe how enzymes sever MTs are illustrated by the “unfoldase” and “wedge” models [Bailey et al., 2016]. The “unfoldase” model is inspired by the canonical mode of action in other AAA+ proteins (such as F1-ATPases and unfoldases). In this model, katanin tears out dimers from the lattice by using the mechanical work of sequential ATP hydrolysis to thread the peptide chain of the tubulin through the central pore of the hexameric severing enzyme complex [White et al., 2007; Johjima et al., 2015; Bailey et al., 2016; Zehr et al., 2017]. As the name suggests, katanin removes subunits from the lattice by pulling away from the surface and unraveling the dimers. Severed tubulin has been shown to reincorporate into nearby MT structures, meaning that the freed tubulin is not likely to be substantially denatured or unfolded [McNally & Vale, 1993]. Alternatively, the “wedge” model proposes that severing enzymes engage the CTT and rigidly hold it within the pore as an anchor while katanin exerts force on the PF interfaces resulting in the destabilization of the lattice and subsequent severing of the structure [Roll-Mecak & Vale, 2008; Bailey et al, 2016; Zehr et al., 2017].

In an effort to decipher the molecular-scale mechanisms of MT severing, we employed the self-organizing polymer (SOP) model [Hyeon et al., 2006] to study the removal of tubulin dimers from a lattice of dimers. In our previous studies, we compared indentation trajectories to AFM studies by applying a force at a constant speed on the MT lattice to simulate the wedge model of severing [Jiang et al., 2017; Szatkowski et al., 2019]. Here, we used SOP to probe the proposed unfoldase mechanism and the possible pathways for filament destruction by severing enzymes. We compared our distributions of MT lattice bending and breaking to experimental results from *in vitro* severing experiments. We employed different 13 PF MT models modeling filaments with different nucleotide states and filament lengths. With our simulations, we can reveal the molecular-scale pathways through which MT lattice break and test their dependence on MT length, accounting for finite size effects. We also investigated the dependence of severing on the location and concentration of severing enzymes on the MT. We find that the experimental data is best fit by a high concentration of severing enzymes working together on three neighboring PFs.

## Results and Discussion

### Pathways for Microtubule Breaking by Pulling on One Subunit

We used simulations to predict the possible molecular scale mechanisms for how MT severing enzymes destroy the MT lattice. In these simulations, we modeled the MT as a cylindrical lattice of dimers, where the interactions between the monomers were extracted from previous molecular dynamics simulations [Kononova et al., 2014]. Such coarse-graining of the interactions allowed us to perform larger-scale simulations of fragments of MTs containing 13 PFs, and up to 16 dimers in length (Fig. S1). We are able to change the geometry of the filament, the interacting substrate, and the mobility of the ends of the filament to try to more accurately capture the situation of real MTs.

We built on our prior simulations using the same simulated system in two different states: the Regular (REG) model, which represents the GDP type lattice, and the All Homogenous model (AHM), which mimics the GMPCPP type lattice [Kononova et al., 2014, Jiang et al., 2017, Szatkowski et al., 2019], as described in the Methods section. Here we examined the unfoldase mechanism where the MT severing enzyme removes dimers through pulling on the CTT(s) of the tubulin dimer. Our approach was to apply a constant loading force on select residues from the CTT end of tubulin monomers (Fig. 1). The orientation of the pulling force was set perpendicular to the long axis of the MT filament. This decision was supported by our studies of pulling at different orientations (see Section SI.1 for details about this choice).

**Figure 1:**
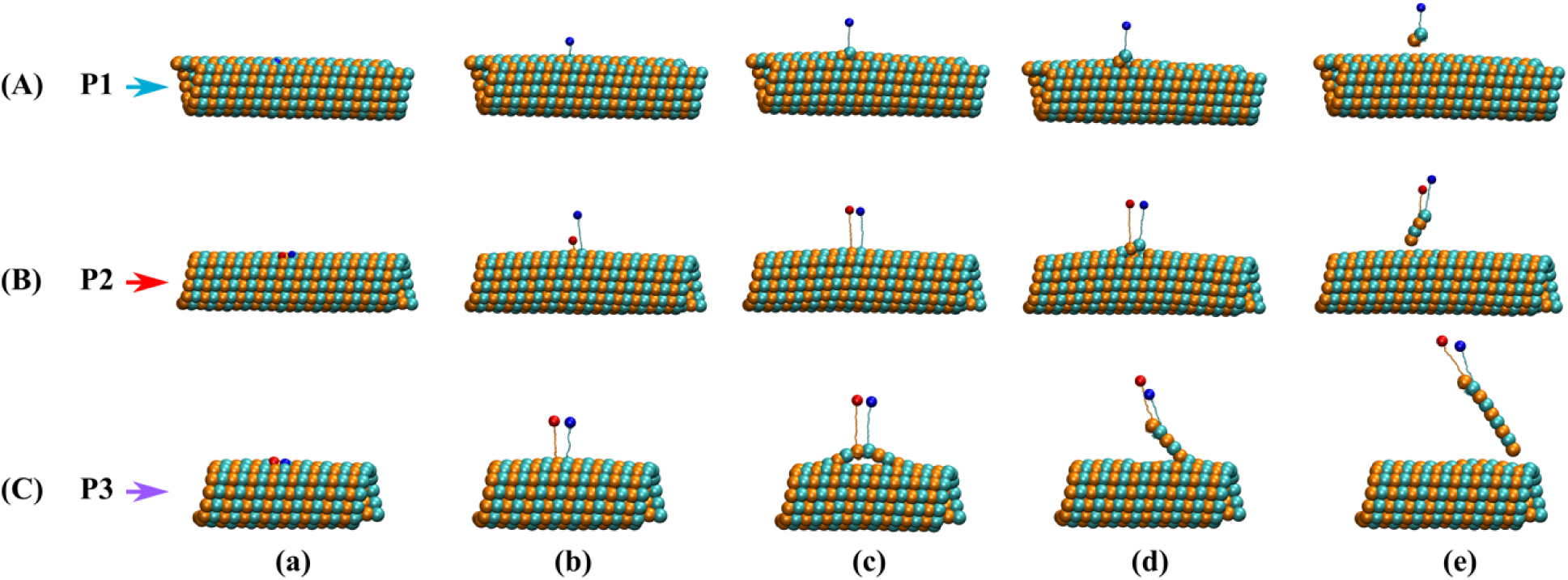
The breaking pathways for MT lattices under pulling forces applied to the C-terminal ends of α (red sphere) and β monomers (blue sphere). (A), (B) and (C) depict the pathways P1, P2, and P3 respectively, described in the text and corresponding to the loss of 1, 2, and respectively, 3 or more dimers from the lattice. For all pathways from left to right we show the conformations corresponding to the: (a) Initial; (b) First breaking force (C-terminal unfolding); (c) Critical breaking force (loss of lateral contacts); (d) Unzipping, and (e) Final state.

In our simulations we found that the MT lattice broke in a variety of ways, which we call pathways. Each simulation can be thought of as a separate experiment and may proceed through a different pathway, as demonstrated in Fig. 1, which shows the most common pathways for MT destruction. The first event in the destruction process, common among all the pathways, is the unfolding of the carboxy-terminal (C-terminal) region. Continued application of force led to the deformation and the breaking of the MT lattice (Fig. 1).

We observed different breaking patterns corresponding to three pathways (referred to as: P1, P2, and P3) leading to the removal of different numbers of dimers from the lattice. The first pathway (P1) is characterized by the loss of lateral contacts between the pulled PF and its adjacent PFs, the loss of longitudinal interfaces that anchor the pulled subunit in the MT lattice, finally leading to the loss of a monomer or a dimer (Fig. 1A). We found this pathway primarily in trajectories involving MTs that are 12 dimers in length, pulling on a subunit located at the seam or when pulling only on the β-terminal end of a tubulin dimer. The second pathway (P2) is only different from P1 in that the lattice eventually loses two dimers (Fig. 1B).

The most complex pathway is the third (P3), mostly observed in shorter MTs of 8 dimers, where we found the loss of lateral contacts between the pulled PF and only one of its neighboring PFs: the PF at its western interface (see depiction of the orientation of interfaces in Fig. S1 in SI). This is followed by the bending of the PF containing the pulled dimer and then, at the critical breaking force, the loss of the lateral contacts between the pulled PF and its other lateral neighbor (the PF on its east side in Fig. S1). Immediately afterwards, one of the longitudinal interfaces made by the pulled dimer with neighboring subunits in the lattice broke and the resulting PF fragment unzipped towards one of the ends of the MT lattice. These trajectories resulted in the loss of more than 2 dimers along a single PF (Fig. 1C). This pathway is particularly exciting, since the removal of long swaths of PFs would result in mechanically unstable filaments, or the ability to create branched MTs, as recently observed in the literature [Basnet et al., 2018].

In all trajectories, the first breaking force or the first event corresponded to the unfolding of the C-terminal regions in the pulled dimer. For the α-tubulin the unfolded region spans helices H-11, H-11’, and H-12 (residues 383-439), while in β tubulin the unfolded region spans helices H-11, H-11’, H-12, and the E-10 β strand (residues 374-429) [Löwe et al., 2001]. The critical breaking force usually corresponded to the loss of contacts between the pulled dimer and one of its lateral neighbors prior to the final longitudinal interface break. In a few cases, such as for MT8 pulling away on the β C-terminal end, the critical breaking force corresponded to further unraveling in the pulled monomer: residues 272 to 373, spanning secondary structure elements H-9, H-9’ to the E-7 β strand.

### Response of MTs to pulling forces applied on a single subunit

#### Simulations for lattices with ends fixed

In our simulations, we have the ability to alter the filament structure and interactions. One particularly interesting avenue to examine was when the small MT segment has free or fixed ends. Fixing the ends more closely mimics a longer MT where the end dimers would be more stable due to interactions with the remainder of the filament, which we do not model. In this set of simulations, we fixed the terminal monomers at both ends of an MT filament and we pulled on either the β-tubulin or on both the α- and β-tubulin C-terminal ends of a single dimer in the center of the outer wall of the MT filament.

Among all 42 trajectories for the MT8 lattice, 17% followed P1 (Fig. 1A) in which the pulled β monomer unfolded up to its E-7 β strand and broke the intra-dimer interface made with the α monomer leading to its removal from the lattice. These simulations corresponded primarily to the pulling only on the β monomer, away from the seam. Another 14% of the trajectories followed P2 (Fig. 1B), leading to a loss of 2 dimers from the lattice, and they correspond mostly to pulling on a β monomer located at the seam. Finally, P3 (Fig. 1C), which was observed in 69% of the trajectories, resulted in the loss of 3 or more dimers and it was the preferred pathway for these simulations. Forces corresponding to the first and critical breaking events for the the MT8 trajectories are listed in Table 1. Representative plots for the evolution of forces along the three pathways for MT8 lattices are depicted in Fig. 2A, while the corresponding plots for all MT8 trajectories are in Fig. S2.

**Table. 1:** Different set-ups, models, average first and critical breaking forces for simulations for pulling on single subunit

| Set-up | MT lattice size,<br>Model | Force (pN) |  |
| --- | --- | --- | --- |
|  |  | First Breaking | Critical Breaking |
| Fixed ends | 8D | $157 \pm 24$ | $361 \pm 41$ |
| | 12D | $210 \pm 32$ | $385 \pm 43$ |
| | 16D | $202 \pm 63$ | $401 \pm 34$ |
| Free plus end | 8D | $205 \pm 76$ | $412 \pm 67$ |
| | 12D | $199 \pm 38$ | $375 \pm 39$ |
| | 16D | $198 \pm 38$ | $377 \pm 44$ |
| Defects | 8D, missing 2 longitudinal dimers, pulling on D44 | $162 \pm 17$ | $507 \pm 15$ |
| | 8D, missing 2 longitudinal dimers, pulling on D57 | $248 \pm 100$ | $298 \pm 48$ |
| | 8D, missing 2 lateral dimers, pulling on D57 | $214 \pm 82$ | $485 \pm 29$ |

**Table. 2:** Different models, average first and critical breaking forces for simulations for pulling on multiple subunits

| Set-up | MT lattice size, Model | Force (pN) |  |
| --- | --- | --- | --- |
|  |  | First Breaking | Critical Breaking |
| **Multi Point pulling** | 12D, Away, 3Pt, Fixed End | 161 | 370 |
|  | 12D, Away, 3Pt, Free Plus End | 140 | 387 |
|  | 16D, Away, 3Pt | 113 | 358 |
|  | 16D, Away, 4Pt (6778) | 197 | 356 |
|  | 16D, Away, 4Pt (6779) | 114 | 206 |
|  | 16D, Away, 4Pt (6789) | 200 | 399 |

**Table 3:** MT lattice models and their respective contact strengths in kcal/mol for intradimer, longitudinal, lateral and at the seam of the lattice

| Model | $\epsilon_h$-value (Contact strength - kcal/mol) | | | |
| --- | --- | --- | --- | --- |
|  | Intra dimer | Longitudinal | Lateral | Seam |
| Regular | 1.9 | 1.0 | 0.9 | 0.9 |
| AHM | 1.9 | average energy per longitudinal interface |  | 0.9 |

**Figure 2:**
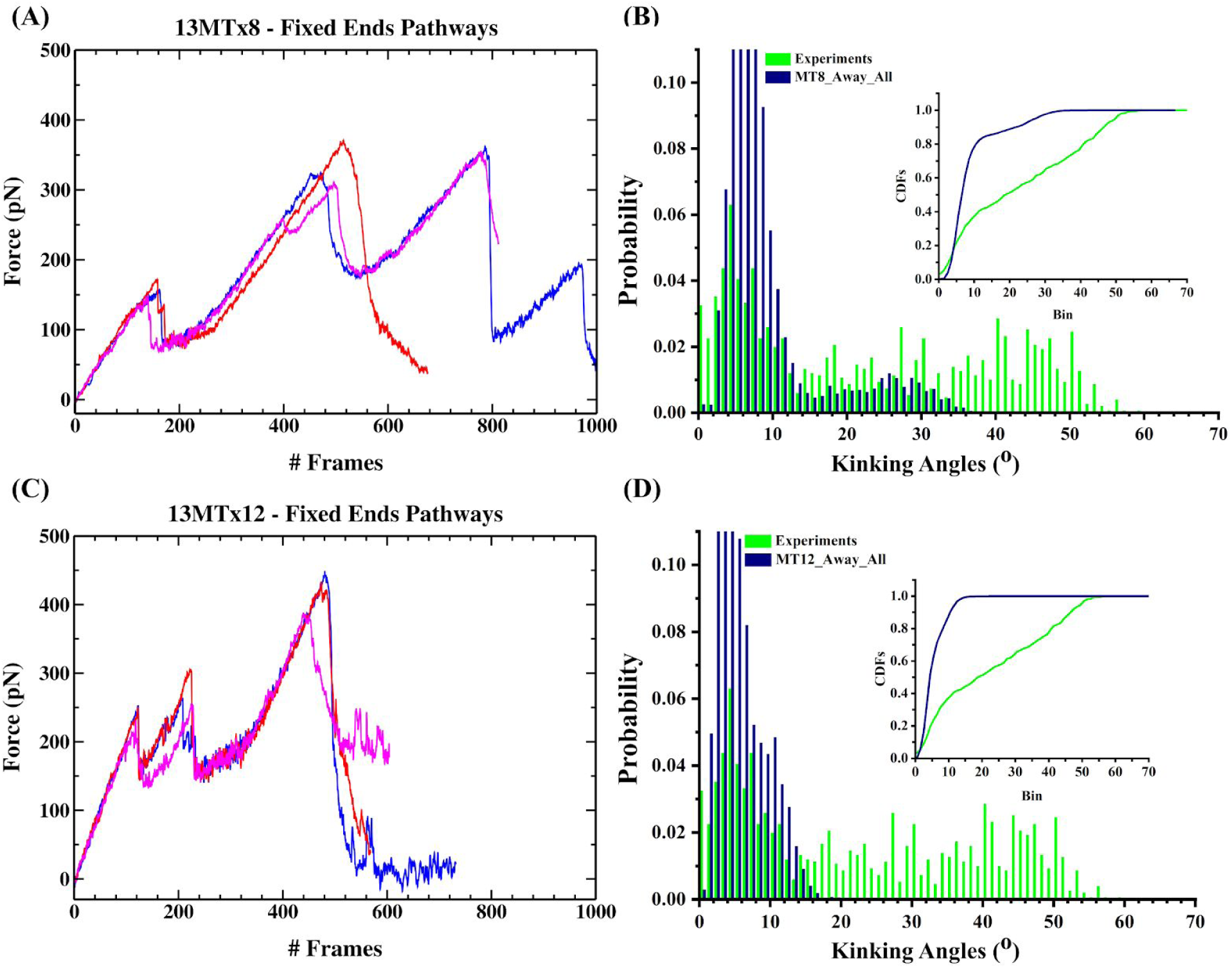
Force vs Frame number (1frame = 0.04 ms) for representative (A) MT8 and (C) MT12 pulling simulations for each of the three breaking pathways: P1 (Blue), P2 (Red) and P3 (Magenta). Distributions of bending angles from (B) MT8 and (D) MT12 pulling away from the seam simulations (blue) and from MT severing assays (Green). The insets show the respective cumulative distribution functions (CDFs).

In order to test our simulations, we have compared the angles of breaking to an experimental realization of MT breaking by severing enzymes. During these experiments, Taxol-stabilized MTs were adhered to a cover slip and the MT severing enzyme, katanin p60, was flowed in while observing in fluorescence microscopy, as described in detail in the Methods. As the MT was severed, free ends of the filament would bend and break. The angles achieved by the MTs during breaking were recorded and histogrammed. For each simulation trajectory, we also calculated the bending angles, using the approach described in the Methods, and compared the simulation histogram to the experimental one (Fig. 2B). The distribution of MT bending angles from our simulations of MT8 filaments showed that most bending angles reside in the 3-12 degrees region, and all were less than 40 degrees (Fig. 2B). This distribution is clearly different from the experimental distribution (Fig. 2B), indicating that the single subunit pulling on the MT8 lattices cannot be used to describe the experimental behavior of MTs during severing. One possible reason for the lack of correlation between simulations and experiments could be the substantial difference in the length of the respective MT filaments (∼ 70 nm in MT8 simulations versus 10 µm in experiments). Although we held the ends fixed to approximate a longer MT, the mobile portion of the filament is shorter, and such finite-size effects are not capable of recapitulating the mechanical response of MTs.

To investigate the possibility that 8 dimers in length is not sufficient, we increased the length of the MT by 50% resulting in a 12 dimers (MT12) long MT structure and by 100% resulting in a 16 dimers (MT16) long MT. For the MT12 lattices, when we pulled a dimer located far from the seam (PF 6), we found an average first breaking force of 209 pN, corresponding to the unfolding of the C-terminal ends of the dimer. When pulling only on β tubulin or on both ends of a dimer, the force for unfolding the β C-terminal region was the same (∼ 166 pN). These results indicate that the higher value of the first breaking force is due to the unfolding of the C-terminal end of the α tubulin monomer. We observed the loss of lateral contacts in the dimer, then between PFs 6 and 7 and PFs 7 and 8 at an average critical breaking force of 398 pN, leading to the loss of a dimer. The behavior corresponded to pathway P1.

When pulling dimers located at the seam, we found a similar series of events: the C-terminal domain unravelled (210 pN), the lateral contacts between PFs 1 and 13 and between PFs 12 and 13 broke (372 pN), a longitudinal interface broke within the pulled PF, and finally, then a second longitudinal interface broke within the pulled PF, which led to the loss of dimer(s) from the lattice. One difference in the behavior of the filament during pulling between the MT8 and MT12 lattices was that the second longitudinal break in MT12 occurred shortly after the first longitudinal break, only allowing for minimal unzipping of the pulled PF. Overall, 68% of the trajectories lost a single dimer (the pulled dimer), following P1. Another 30% of the trajectories followed P2 by losing 2 dimers, which was the primary pathway in simulations that involved the AHM lattice and pulled on both C-terminal ends of the central dimer. The third pathway (P3) was found in only 1 trajectory and was characterized by the loss of 5 dimers. The average forces obtained for the three pathways in MT12 are shown in Table 1 and examples of the force evolution along each pathway are in Fig. 2C. Details for all the simulations are in Table S1 and in Fig. S3. An important outcome from these simulations is that MT12 lattices have a lower propensity to lose a large number of dimers compared the shorter MT8 filaments, which is indicative of finite-size effects in the mechanical response of the MT8 lattices.

We compared the bending angles of the MTs in the simulation with experimental results from the MT severing assays. The distribution of the bending angles from the MT12 simulations is populated only in the 3-18 degrees region. The MT12 results are even more contrasting from the experimental distribution than the MT8 simulations (Fig. 2D). Upon increasing the size of lattice to 16 dimers i.e., to twice the size of MT8, we found only two of the three breaking pathways discussed before: P1 when we pulled only on the β C-terminal end and P2 when we pulled on both C-terminal ends (Fig. S4). Again, as discussed above, this is a manifestation of finite-size effects. The breaking forces for these pathways were similar to those found in the MT12 simulations. Moreover, the bending angle distribution was similar to that for the MT12 case (Fig. S3E and S3F) and thus not a good fit for the experimental distribution.

In summary, our simulations strikingly show that, unlike the MT lattice deformation under the action of indentation forces [Szatkowski et al., 2019], the correlation between the distributions for bending angles from increasingly longer lattices (from MT8 to MT16) and the experimental distribution decreases dramatically. An increase in the length of the filament should bring the simulated MTs closer to experiments as we observed with indentation. Thus, the opposite behavior of filaments under the action of forces localized on a subunit rules out pulling on a single subunit in the middle part of an MT filament (away from the ends) as a severing mechanism.

#### Simulations for lattices with the Plus End free

In our simulations, we were able to alter the forces that the simulated MT experienced to test how different circumstances affect the pathways to removing dimers. Here, we fixed only the minus end of a MT lattice, keeping the plus end free. This set-up allowed us to test the role played by freely-fluctuating ends in the response of MTs to pulling forces applied close to such ends. We found that the physical changes in the MT8 lattice when pulling on a single dimer, on and away from the seam, were similar to the pathways observed when both ends of the lattice were kept fixed. All trajectories involving pulling only on the β C-terminal end, away from the seam (Fig. S5A), started with the unfolding of the dimer (177 pN), followed by the loss of lateral contacts between the pulled PF and one of its neighboring PFs (336 pN), and the loss of a longitudinal interface that led to the unzipping of the pulled PF towards the minus end. The critical breaking force (356 pN) corresponded to further unfolding of the β monomer and the loss of the pulled monomer, according to P1.

When pulling on both the α and β C-terminal ends of the same dimer (Fig. S5C), the first breaking force (214 pN) corresponded to the unfolding of the α-C-terminal end, followed by the unfolding of the β C-terminal end. The critical breaking force (375 pN) coincided with the loss of contacts between the pulled dimer and one of its lateral neighbors in the lattice. This was followed by the loss of a longitudinal interface in the pulled PF and the unzipping of the resulting PF fragment towards the plus end, losing more than 2 dimers, according to P3.

The force required to unfold the C-terminal end, as well as the typical critical breaking force, for pulling on the MT8 lattice with the plus end free were ∼ 50 pN higher than in the fixed ends simulations. We found that this is the consequence of a shift in pathway population for MT8: from most of the fixed ends trajectories following P3 to only a minority of the free plus end trajectories doing so. This change, combined with the fact that both the first and critical breaking forces for P3 are lower by ∼ 50 pN compared to the values for P1 and P2, account for the increase in the average force values for the free plus end simulations of MT8 lattices.

MT destruction pathways for simulated MT12 and MT16 filaments correspond to the loss of at most 2 dimers - no matter the method for pulling. Comparing the critical force values for MT8, MT12, and MT16 filaments for both fixed and free ends (Table 1), we observe that the MT8 filaments match those for the longer MT12 and MT16 filaments, which indicates that the release of constraints at one end of the lattice minimizes the finite-size effects of pulling on single subunits from short MT lattices. Comparing the experimental bending angles to the simulation results for MT8 with one free end, the distribution of bending angles from the plus end free simulations is even farther away than the one for both ends fixed from the distribution found in severing experiments, indicating that the plus end free set-up does not account for MT severing (Fig. 3B).

**Figure 3:**
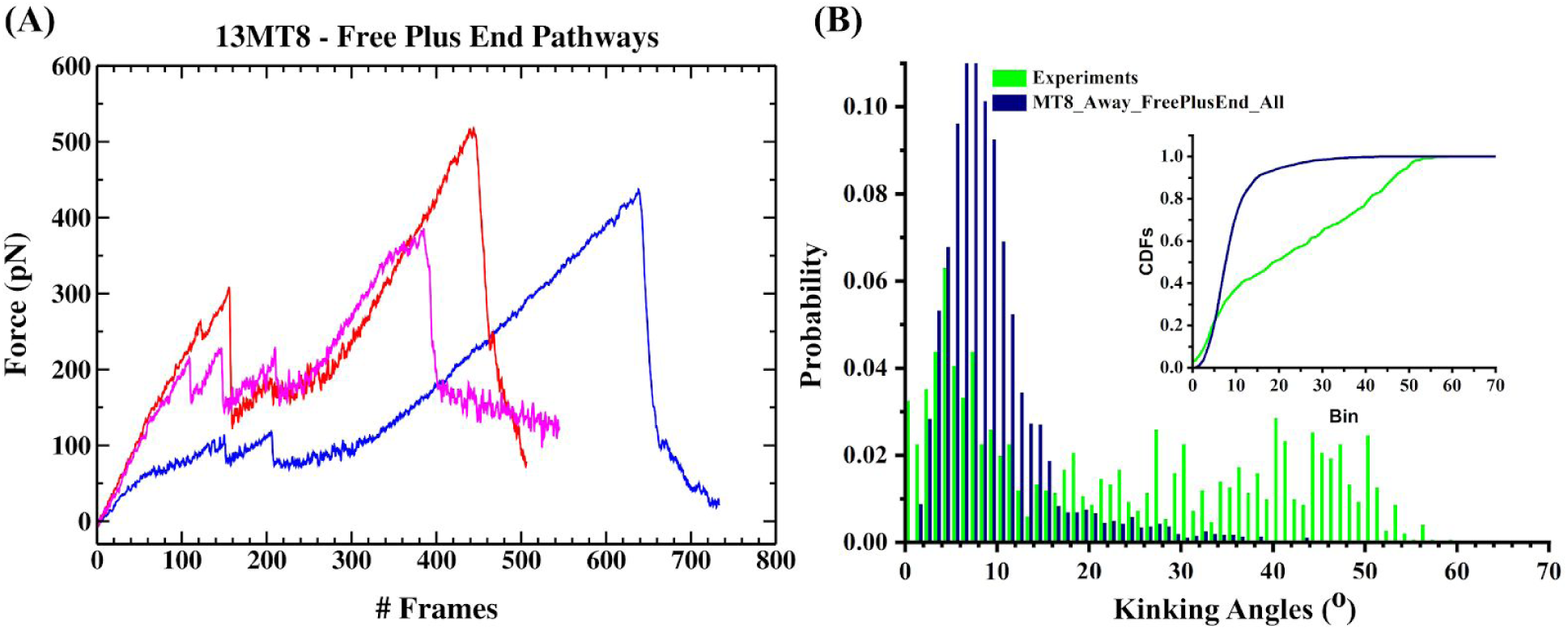
(A) Force vs Frame number (1 frame = 0.04 ms) for representative runs of pulling away on MT8 with the plus end free. The runs represent the three breaking pathways: P1 (Blue), P2 (Red) and P3 (Magenta). (B) Distributions of bending angles from MT8 pulling away (blue) and from MT severing assays (Green). The insets show the respective cumulative distribution functions (CDFs).

#### Simulations for lattices with defects

Following reports from the literature, we previously probed the response of MT lattices with missing subunits to mechanical forces applied akin to the set-up from AFM indentation experiments [Jiang et al., 2017]. In that case, we found that a MT lattice with 2% defects i.e., missing either 2 longitudinal or lateral dimers, had a weakened mechanical response to indentation forces compared to intact lattices. To test whether the defects are relevant for the response of a filament to pulling forces applied to individual subunits, we carried out simulations by pulling on the dimer next to a lattice defect such that the dimer being pulled had a missing lateral interface (Fig. 4).

**Figure 4:**
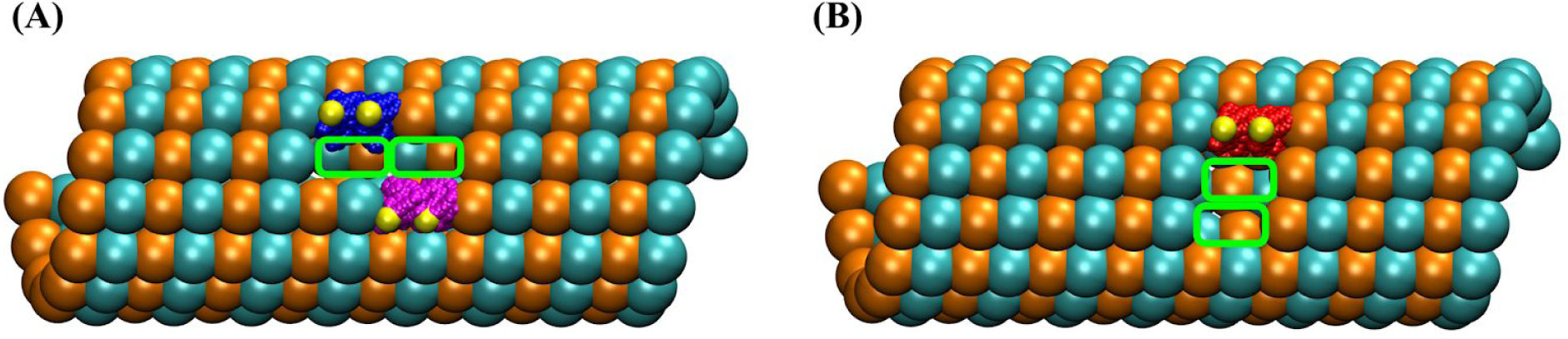
Representation of 8 dimers long MT with 2% defects (A) with 2 longitudinal dimers (D45 and D58) missing pulled on adjacent D44 (Blue) or D57 (Magenta) dimers. (B) with 2 lateral dimers (D58 and D59) missing pulled on D57 (red). Yellow: Pulled C-terminal region residues on the respective dimers.

The breaking pathways for these simulations resemble those articulated for defect-free MTs, but there are some important differences. For instance, while the first breaking event for pulling only on the β C-terminal end of a subunit corresponded to β C-terminal unfolding (same as in the defect-free case), in trajectories where we pulled on both C-terminal ends of a subunit the first event was the loss of lateral contacts between PFs 5 and 6 leading to the formation of a crack that propagated towards the fixed ends of the lattice (Figs. S8 and S10), followed by the unfolding of the two ends. This release of the lateral contacts between PFs 5 and 6 allowed the pulled PF to collapse on its other neighboring PF resulting in a delay in the breaking of the filament and in large changes to the values of the critical breaking forces compared to the defect free MTs.

Our simulations of MT lattices with 2% defects were performed using three different set-ups: (1) lattice with 2 longitudinal dimers missing in PF 6 (D45 and D58) and pulled on D44 in PF 5 (blue in Fig. 4A) which have no eastern lateral neighbor, (2) lattice with 2 longitudinal dimers missing in PF 6 (D45 and D58) and pulled on D57 in PF 7 (magenta in Fig. 4A) which have no western lateral neighbor, and (3) lattice with 2 lateral dimers missing (D58 and D59) in PF 7 and PF 8, and pulled on D57 in PF 5, located next to the missing subunits (Fig. 4B). The notation for the missing and, respectively, the pulled dimers follows Fig. S1.

Among the 31 simulated pulling trajectories, 10% followed pathway P1′ (Fig. S8) where the initial β C-terminal unfolding occurred at a force of 167 pN, followed by the breaking of a longitudinal interface and finally the loss of the pulled dimer. The second pathway (P2′, Fig. S9), which is the most probable (55% of the trajectories), corresponded to the loss of the lateral contacts between PFs 5 and 6 near the defects resulting in crack propagation. The next event, which was the unfolding of the C-terminal end, was followed by a first longitudinal break and the unzipping of the resulting fragment of the pulled PF towards the minus end. Finally, the lattice lost 2 dimers. The third pathway (P3′, Fig. S10), found in 35% of the trajectories, was similar to P2′ but in this case the PF lost more than 2 dimers.

The critical breaking forces for pulling the C-terminal ends of dimers according to set-ups (1) and (3) are 507 pN (Figs. S6A and S6D) and 485 pN (Figs. S6C and S6F), respectively (Table 1). These forces account for the break of the lateral contacts between PFs 4 and 5 and the break of two longitudinal interfaces in PF 5, leading to the extraction of a fragment from this PF (Fig. S7B). This catastrophic breakdown of the contacts between the pulled region and the rest of the MT lattice gives rise to unusually large critical breaking forces (Fig. S7B) and shows that the MT8 lattice becomes very brittle compared to the defect-free case, described above, where the lateral and longitudinal breaks occur independent of each other (Fig. S7A). In contrast, set-up (2) led to a critical breaking force of only 297 pN (Figs. S6B and S6E), which was ∼ 100 pN lower than the force for the defect-free MT (Table 1). For this configuration, the critical breaking force accounts only for the bending and breaking of the pulled PF, i.e., of PF 7. In addition, when we pulled on both C-terminal ends, we found that only the β C-terminal unfolds.

Interestingly, we found that some of the bending angle distributions resulting from these simulations are distinct from the distributions obtained when pulling on an intact MT lattice. In comparing these distributions with the kinking angle distributions from experimental *in-vitro* severing assays, the behavior of the MT lattices with defects from set-ups (1) and (3) was more similar to experiments than our simulations intact lattices (Figs. S2E and S2F). In contrast, simulations of a MT lattice with defects pulled according to set-up (2) led to similar bending angle distribution (Fig. S6H) to that found for the simulations of defect-free lattices with fixed ends (Fig. S2E); both were different from experiments.

We found that the two contrasting behaviors for the bending angles in simulations result from the degree of collective bending of sets of PFs (Fig. S7C and S7D). Specifically, the bending angles for both the pulled PF and the PF at its eastern interface are high and track each other all the way up to the point of the breakdown of the pulled PF, when pulling on subunits with missing neighbors (Fig. S7D). In contrast, for the defect-free intact MT8 lattice, the degree of bending in the pulled PF is consistently low throughout the trajectory and none of its neighboring PFs exhibit any pronounced bending (Fig. S7C). Moreover, we found that only the simulations that result in the bending and breaking of multiple PFs, versus that of a single PF, led to bending angle distributions (Figs. S6G and Fig. S6I) resembling the *in vitro* severing distribution.

### Response of MTs to pulling forces applied on multiple subunits

So far, we have tested the contributions of pulling on the C-terminal ends of a single subunit to create lattice breaks, and found that most do not resemble the experimental results for severing. This outcome is in stark contrast with the results of our previous MT indentation studies, which led to good or excellent matches with the bending angle distributions of MTs in severing experiments [Jiang et al., 2017; Szatkowski et al., 2019].

A clear difference between the behavior of the MT lattice during indentation versus pulling on a single subunit is that indentation always leads to the bending and breaking of multiple PFs. In contrast, during single subunit pulling, usually only the pulled PF undergoes bending and breaking. The only exceptions are for certain situations when lattices have missing subunits, where we found cooperative bending and breaking of two neighboring PFs resulting in bending angle distributions that come closer to experimental results. The difference in outcomes between the two manners of force application suggests that cooperativity in PF breaking is significant for severing.

To ascertain the level of cooperativity in PF bending and breaking needed to account for experimental findings during severing, we carried out new simulations where we pulled, at the same time, on the C-terminal ends of multiple dimers in a MT lattice. We used 12 and 16 dimers long lattices in this set of simulations, as well as different numbers of pulled dimers per lattice length. Operationally, we pulled on 3 dimers from the MT12 and MT16 lattices and also on 4 dimers for the 16 dimers long MTs. In all cases, we selected dimers on neighboring PFs located at positions distant enough to accommodate the placement of the full oligomeric (hexameric) state of the severing enzymes without steric clashes (Fig. S11). In these studies, changing the number of dimers being pulled gives insight into the concentration dependency of the action of severing enzymes.

Following our above approach for the single-point pulling runs, we classified the multi-point pulling breaking patterns into different pathways (MP-1, MP-2 and MP-3) based on how the PFs bend, lose dimer(s) and break from the MT lattice (Fig. 5). For all three pathways, the first event corresponded to the unfolding of the C-terminal ends of the pulled dimers, which is the same behavior we found in the three pathways for the single subunit pulling reviewed previously (Fig. 1). The next steps differentiate between pathways. In the MP-1 pathway, pulling on multiple dimers led to the fast and concerted loss of lateral and longitudinal contacts resulting in the removal of a single dimer from the lattice. We observed this pathway for the MT16 lattice when pulling on PFs 6, 7, 7 and 9 (Fig. 5 (MP-1) and Fig.S11F). The second pathway (MP-2) is characterized by the loss of lateral contacts resulting in the formation and propagation of cracks towards the ends of the lattice, and then the removal of dimers from PF 7. Next, we found longitudinal interface breaks and substantial bending of the pulled PFs resulting in the loss of the pulled dimers. We observed this pathway in simulations of the MT12 and MT16 lattices, when pulling on 3 dimers (Fig. 5 (MP-2), and Figs. S11C and S11D). The final pathway (MP-3), found for the 4 point pulling on PFs 6, 7, 7 and 8, and on PFs 6, 7, 8 and 9 of the MT16 lattice (Fig. 5 (MP-3), and Figs. S11E and S11G), resulted in the loss of the lateral interfaces between PFs 5 and 6, and PFs 9 and 10 forming a crack that propagated towards the ends of the filament. Then the lattices lost two dimers and the longitudinal interfaces formed by the pulled PFs with the fixed end dimers at the minus end of the lattice (Fig. 5D (MP-3)). A common denominator of pathways MP-2 and MP-3, which distinguishes them from pathways P1 to P3 from the single subunit pulling, is that in both cases groups of neighboring PFs undergo slow correlated changes such as bending, and longitudinal breaking. More information regarding the pathways for various setups is provided below, with full details in the Section SI.2.

**Figure 5:**
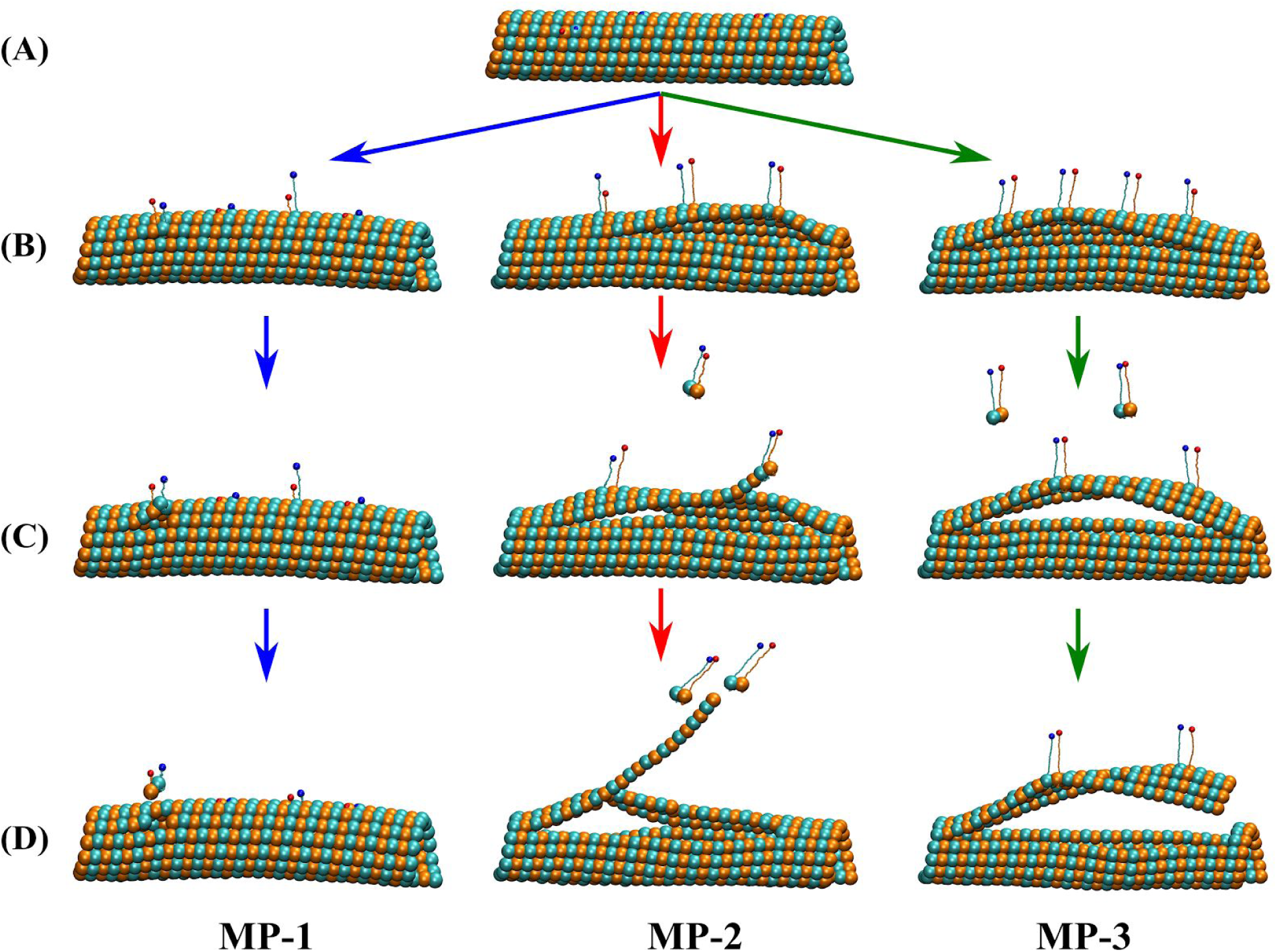
Breaking pathways for MT lattices under pulling forces applied on multiple positions at the C-terminal ends of α (red sphere) and β monomers (blue sphere). MP-1, MP-2 and MP-3 indicate different pathways. For all pathways from top to bottom we depict the conformations corresponding to the: (A) Initial; (B) C-terminal unfolding and loss of lateral contacts; (C) Critical breaking force; and (D) Final state.

#### Pulling on 3 dimers from 12 dimers long MTs

As described above, we applied pulling forces on MT12 lattices at 3 locations chosen such as three severing enzymes could fit close together on the lattice (Fig. S11C). We first simulated filaments with the ends fixed. We pulled on three dimers from three different PFs: PF 6 (D110), PF 7 (D72) and PF 8 (D34) (Fig. S12). The breaking of the lattice follows pathway MP-2 (Fig. 5). The first event, the unfolding of the C-terminal ends of the dimers closest to the fixed ends, happened at a force of 161 pN per pulled position. The break of the longitudinal interfaces at the fixed plus ends of PFs 7, 8 and 9 corresponded to the critical breaking force of 370 pN. The final event was the loss of 2 dimers from PF 8 (Fig. S12F). The bent fragment from PF 6, seen in Fig. S12F, will likely detach from the MT lattice, but the time for this event exceeds the 50 ms run time in our simulation.

Next, we examined the same MT12 filaments with the plus end free. We selected the same 3 subunits on PF 6 (D110), PF 7 (D72) and PF 8 (D34) for pulling (Fig. S13). In this set of simulations, the first event corresponded to the β C-terminal unfolding of the dimer closest to the fixed (minus) end at a 140 pN force (Fig. S13B). This is in contrast to the simulations with both ends fixed where the first event was the unfolding of the C-terminal ends of all the pulled dimers. For the free plus end simulations, the critical breaking force of 387 pN corresponded to the loss of lateral contacts between PFs 8 and 9, and the loss of 4 and 2 dimers from the peeled off fragments of PF 6 and PF 8, respectively (Fig. S13F).

The simulations with the ends of the lattice kept fixed result in higher bending angles than the simulations with one end free. When comparing the bending angle distributions from these two simulation arrangements with the experimental angles (Figs. 7A and 7B), we found that the simulations of MT12 with fixed ends are the better fit to the experimental data.

**Figure 6:**
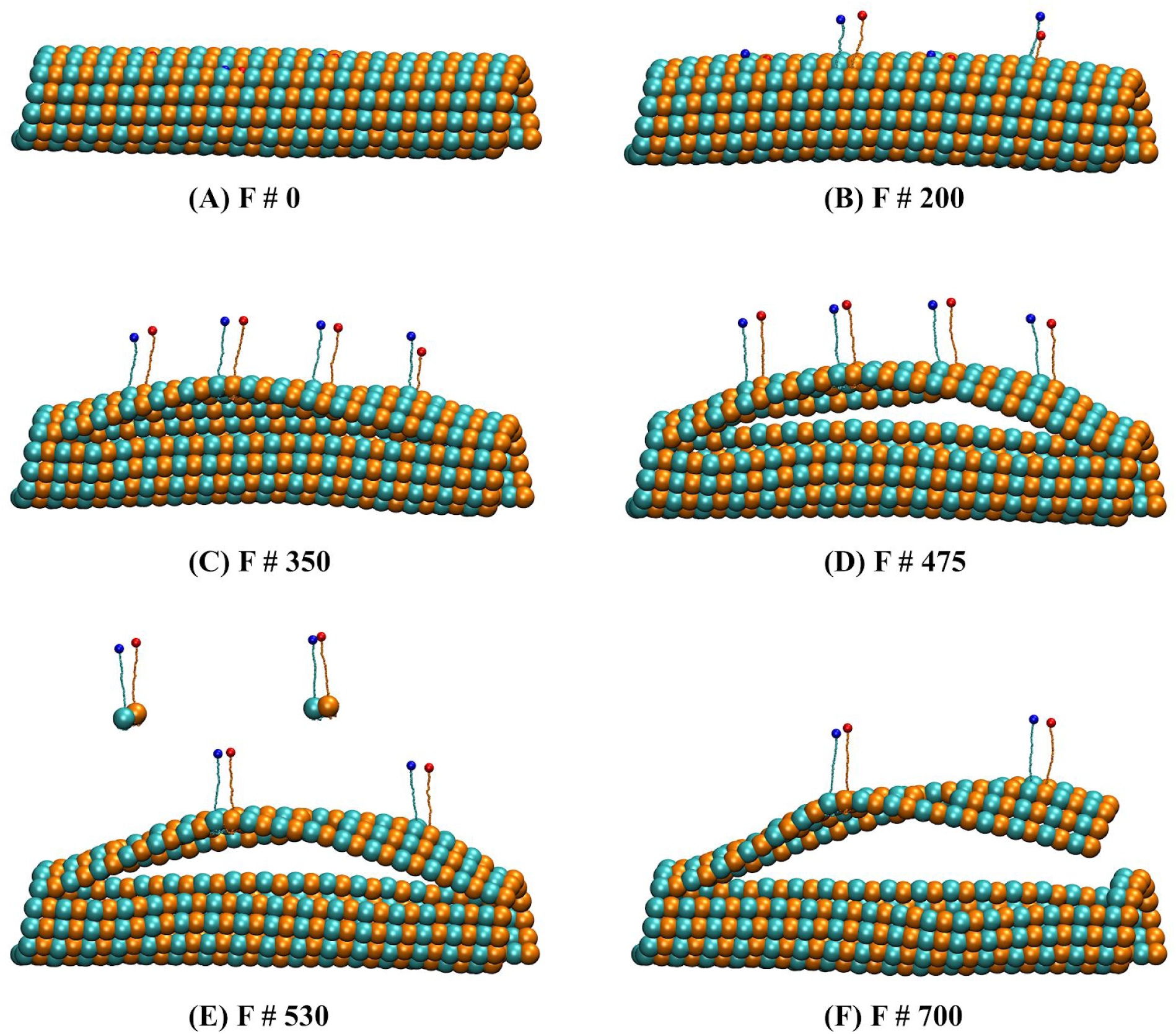
Representative states for the 4 point pulling on PFs 6, 7, 7 and 8 of a MT16 lattice. (A to F) showing different states (F # - frame number; 1 frame = 0.04 ms) along the trajectory. We pulled on β (Blue) and α (Red) C-terminal ends of dimers D47, D85, D123 and D163.

**Figure 7:**
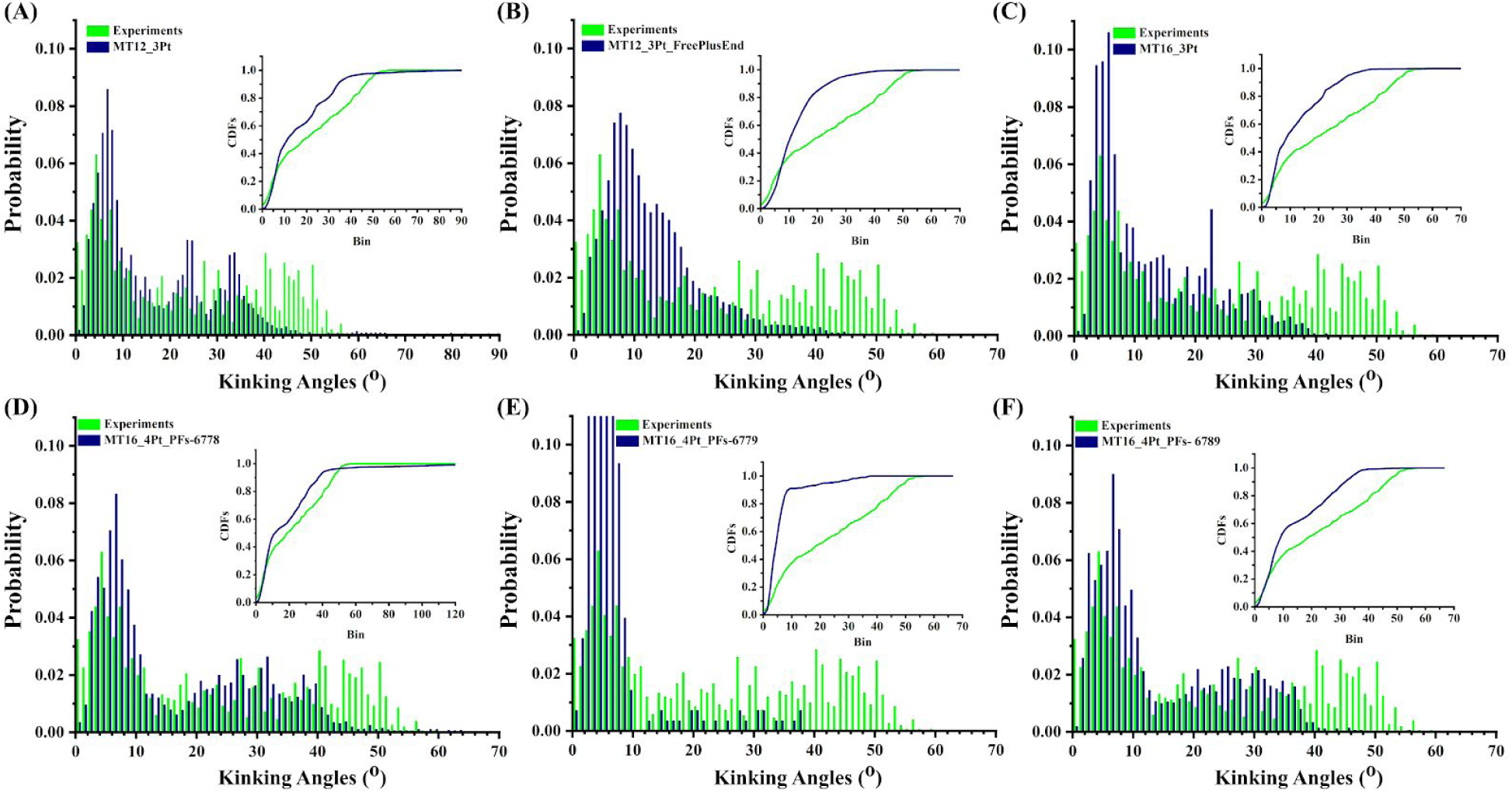
Distribution of bending angles from severing assays (Green) compared to simulations (blue) of 3 point pulling on (A) MT12 with fixed ends, (B) MT12 with free plus ends, (C) MT16 with fixed ends, and 4 point pulling on (D) MT16 pulled on PFs 6,7,7, and 8, (E) MT16 pulled on PFs 6,7,7, and 9, (F) MT16 pulled on PFs 6,7,8 and 9.

#### Pulling on 3 dimers from 16 dimers long MT

The purpose of simulating long filaments was to more closely approximate real MTs. In our 3 point pulling on the MT12 lattice, we found the bending behavior from the fixed ends series most closely resembled the experimental data, so we only used fixed ends for the simulations of MT16 lattices. We pulled on 3 dimers from 3 different PFs (Fig. S11D): PF 6 (D45), PF 7 (D98), and PF 8 (D151) (Fig. S14). As discussed above, the breaking of the lattice follows pathway MP-2 (Fig. 5). The first breaking force of 113 pN corresponded to the unfolding of the β C-terminal end of D151. The loss of the lateral interface between PFs 9 and 10 led to the formation of a crack that propagated towards the minus end and to the loss of D98 from PF 7 at a critical breaking force of 345 pN (Figs. S14D and S14E).

We found that the first and the critical breaking forces for the 3 point pulling decreased with the length of the MT filament (Table. 2). Moreover, while in the MT12 simulations the first breaking event corresponded to the unfolding of both C-terminal ends of the pulled dimers, in the MT16 case the first event corresponded only to the unfolding of the β C-terminal end of one dimer. Thus, the unraveling events were more localized for longer MT lattices. Importantly, the fact that the pulled dimers were located closer to the fixed ends of the filament in the MT12 case resulted in the increase in force and in the removal of a higher number dimers from the shorter lattice. In turn, this meant the bending angle distribution from MT12 simulations are closer to the experiments (Fig. 7A) than MT16 simulations (Fig. 7C).

#### Pulling on 4 dimers from 3PFs of the 16 dimers long MT

Our simulations for pulling on 3 dimers from MT lattices of two different lengths showed that, if the pulled subunits are located closer to the ends of the filament the bending and breaking pattern came closer to the experimental behavior. To further investigate this point, we carried out pulling studies on MT16 lattices under conditions mimicking higher severing enzyme concentration: we applied pulling forces on 4 subunits (Fig. S11).

We pulled on 4 dimers from 3 PFs (Fig. S11E): PF 6 (D123), PF 7 (D85 and D163), and PF 8 (D47) mimicking the addition of another severing motor on the lattice such that 2 motors act on subunits from the same PF. The changes in the MT lattice in this set-up followed the pathway MP-3 (Fig. 6). The initial breaking event at a 197 pN force corresponded to the unfolding of C-terminal ends in the dimers closest to the fixed ends of the filament (Fig. 6B). At the critical breaking force of 356 pN, the pulled PFs broke from their longitudinal interfaces with the fixed dimers at the minus end of the lattice (Fig. 6F) and unzipped towards the plus end. When comparing the distribution of bending angles from this type of simulation and severing experiments, we observed a good agreement (Fig. 7D). The computational distribution is bimodal, consisting of one highly populated peak at low angles and a second peak at higher angles reaching up to 50 degrees.

To evaluate the role played by the correlated response of PFs in the bending and breaking of MT lattices under the action of pulling forces, we kept the concentration of the modeled enzymes intact, but instead of pulling on 4 dimers located on 3 consecutive PFs, we pulled on dimers from non-neighboring PFs (Fig. S11F), i.e., from PFs 6, 7 and 9 (D48, D85, D123 and D163). These simulations followed pathway MP-1 (Fig. 5). The first breaking event at 114 pN was due to the C-terminal of dimers on PF 6 (D85) and PF 9 (D163) unfolding (Fig. S15D). The critical breaking force at 206 pN corresponded to the unraveling of dimers on PFs 6 and 9, the loss of lateral contacts between D48 and D49, and the partial unfolding of the β C-terminal end of D85 from PF 7 (Fig. S15E). The bending angles distribution (Fig. 7E) is similar to the one for the single point pulling on lattices with ends fixed, thus being a very poor fit for the experimental distribution. This finding, combined with our results for pulling on positions from PFs 6, 7, and 8, shows that the presence of strong correlated changes in the pulled PFs is a requirement during severing.

#### Pulling on 4 dimers from 4PFs of 16 dimers long MT filaments

Because pulling on multiple dimers located on 3 neighboring (consecutive) PFs led to better matches with the experimental data than pulling on subunits from non-adjacent PFs, we next explored the role played by the number of consecutive pulled PFs in rendering behavior that can recapitulate the experimental bending angle distribution. This time, we pulled on 4 dimers (D47, D85, D123 and D165) from 4 consecutive PFs (Fig. S11G): PFs 6 to 9 (Fig.S16). These simulations followed pathway MP-3 (Fig. 5). The unfolding of the C-terminal ends of the pulled dimers on PFs 6, 8 and, respectively, 9 resulted in the first breaking event at 200 pN (Fig. S16B). The critical breaking force of 400 pN led to further unraveling of the C-terminal ends of some of the pulled dimers (Fig. S16E), to the break of the longitudinal interfaces of all pulled PFs from the fixed minus end, and the unzipping of the resulting set of PF fragments towards the plus end (Fig. S16F). The resulting bending angle distribution (Fig. 7F) reproduced some aspects of the experimental distribution, but it was not as good a fit as when we pulled on 4 dimers from the 3 neighboring PFs 6, 7, and 8 (Fig. 7D). These findings indicated that, while the increase in the number of pulled consecutive PFs, which is a stand-in for an increase in the concentration of severing enzymes, is important for severing if it occurs by an unfoldase mechanism, the most important factor for achieving a good match between simulations and experiments would be the maintenance of a high degree of correlated changes in the pulled PFs for as long as possible before the extraction of fragments from the lattice. Thus, both the concentration and the placement of severing enzymes on an MT lattice are important for efficient severing.

The analysis of multipoint pulling simulations for MT12 and MT16 lattices with fixed ends detailed in Section SI.3, showed that pulling on multiple PFs led to pronounced permanent bending in other PFs (Fig. S18). We expect that this bending will speed up the breaking of additional PFs in subsequent severing steps.

Our work provides new insights into the mechanism of MT severing: while the interaction between a single MT subunit and the hexameric state of the severing enzyme is a requirement for the binding/attachment of the enzyme on the surface of the MT, the application of a pulling force on only one dimer at a time does not reproduce the experimental results from *in vitro* severing assays. In contrast, the coordinated action of multiple enzymes of pulling on dimers from a number of PFs, which are lateral neighbors in the MT lattice, is a more likely match for the severing experiments. This finding, combined with our previous studies of MT lattice indentation [Jiang, 2017; Szatkowski et al., 2019], offers strong support for a model where severing is achieved by the coordinated action of multiple copies of a severing enzyme: each enzyme applies a pulling force on the tails of a tubulin dimer, resulting in the extension and likely translocation of the tail inside the pore of the enzyme. The reaction to this upward force results in a wedging force being applied on the MT surface akin to indentation, which is an efficient manner for breaking interfaces between subunits leading to the fragmentation of a number of PFs; continuous application of the pulling forces on dimers from the broken PF fragments eventually leads to extraction of subunits from the filament. Future simulations could investigate combined pulling and pushing schemes, as MT severing enzymes could be applying localized forces in both directions at the same time.

## Methods

### Computational Methods

#### Total Potential Energy for the SOP Model

All simulations were performed using Brownian dynamics and the self-organized polymer (SOP) model accelerated on GPUs (gSOP version 2.0) [Hyeon et al., 2006a]. The model uses the equation seen below to determine the different interactions within the protein, described by the total potential (V_**T**_), that will dictate the behavior of the structure in response to specified forces on the lattice. The finite extensible nonlinear elastic (V_**FENE**_) potential represents the backbone of the structure, the full Lennard-Jones potential (V^**ATT**^_**NB**_) represents the native non-bonded interactions in the structure, while the repulsive Lennard-Jones potential (V^**REP**^_**NB**_) represents the non-native non-bonded interactions in the structure:

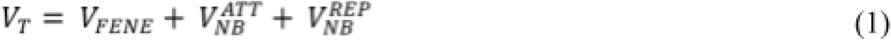

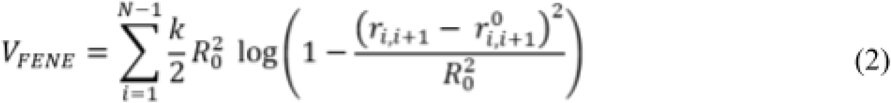

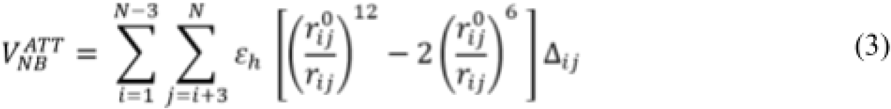

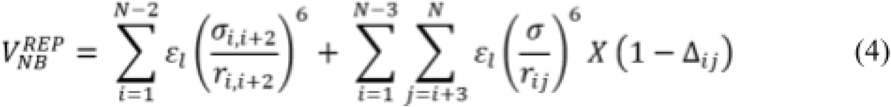

The strength of contacts for the non-bonded native interactions is given by the parameter ε_h_ in eq.3, which changes based on the lattice model used. The values for the contact strength (ε_h_) for each of the two MT models used in our simulations are listed in Table. 3. The remaining parameters are: the frictional coefficient (ζ) set to 50, the spring constant (k) for covalent interactions set to 20.0 kcal/mol·Å, R_0_ = 2.0 Å, and r_i_ = 3.8 Å, for i = 1, 2, …, N, where N is the total number of residues. r_i,j_ represents the distance between two residues, i and j, while r_i,j_^0^ is its value in the native structure. The other parameters specifically used in pulling simulations are the cantilever spring constant, k_trans = 0.05 kcal/mol^2^, deltax (Δ*x*) = 0.0008 Å, i.e, the displacement of the cantilever during a simulation from which the pulling speed (***v***_***f***_) is calculated using Δ*x*/(n_*av*_*\** Δt). We performed our simulations by pulling on selected dimers, as described in the text, at a speed of 2 μm/s using a temperature of 300 K. Because the integration time step was 40 ps, every frame in our plots corresponds to 40 μs. As mentioned above, in our simulations we used 13 PFs MT lattices with two different topologies: the Regular (REG) model, which represents the GDP type lattice, and the All homogenous model (AHM), which mimics the GMPCPP type lattice, as discussed in our previous work [Jiang et al., 2017]. To evaluate the influence of the finite size of the MT filament effects on the results, we carried out our studies on lattices of increasing length: 8, 12, and 16 dimers long. The orientation, numbering of the PFs and the dimers can be found in Fig. S1.

#### Simulations Setups and Data Analysis

Due to the lack of density for the CTTs in the α-β tubulin crystal structure (PDB ID: 1JFF) [Löwe et al., 2001] used in construction of MT lattice for our simulations, we pulled on the terminal residues with actual density in the C-terminal region of the α- and β-tubulin monomers. We applied point forces on selected residue(s) from the C-terminal regions of β-tubulin (β) and from α- and β-tubulin (both) at a speed of 2.0 µm/s, the same speed used in our indentation simulations as well as in AFM experiments [Szatkowski et al., 2019]. We selected dimers located either on the seam (the A-lattice region of the MT filament) or away from the seam (the B-lattice region of the MT filament) [McIntosh et al., 2009]. We used four set-ups: (i) Pull on a single dimer when both ends of the MT lattice are fixed (plus and minus end), (ii) Pull on a single dimer when the plus end of the MT lattice is free to fluctuate (Free plus end), (iii) Pull on dimer(s) from lattices with missing subunits and, (iv) Pulling on multiple dimers from an MT lattice (Fig. S1). We selected the direction of pulling as perpendicular to the long axis (Z-axis) of the MT lattice and away from the outer surface of the filament. We ran Brownian dynamics using the same parameters used in our previous work with an integration time step of 40 ps [Jiang et al., 2017].

#### Data Analysis

For each of the above-listed set-ups we ran multiple trajectories and analyzed the resulting force-extension curves, Q_n_ (percent of native contacts during the simulation), and the evolution of the bending angles. We compared our distributions with the distribution of kinking angles obtained from *in vitro* severing experiments. We categorized all the simulations based on their common pathways.

#### Force-extension/distance Curves and Q_n_

From each simulation, we obtained the force response curve and we used it to identify the forces corresponding to the C-terminal unfolding, the initial lattice breaking, the first longitudinal breaking and the critical breaking. The idea is that such sets of breaking events would determine how severing enzymes may cut MTs when employing an unfoldase action. The force response curves varied depending on the breaking pathway. The forces for each breaking event for individual trajectories and for each setup are listed in Table. T1 and Figs. S2 - S6, S17 and S19B. To determine whether the lattice lost an interface or not, and to determine the extent of the interface fracture during the simulation, we monitored the time evolution of the native contacts in the lattice. At each frame we used a tolerance of up to 13 Å above which the attractive potential (Lennard-Jones) in Eqn.(1) becomes insignificant, to denote that a pair of residues still forms a native contact. We applied this type of analysis for the intra-dimer contacts between α and β tubulin interface in a given dimer, and for the inter-dimer contacts formed between two neighboring dimers. We further categorized the inter-dimer contacts into north (N), south (S), east (E), or west (W) based on the interface they shared with the pulled dimer(s) (depicted in Fig. S1). Namely, for any dimer located away from the ends of the MT filament, there are contacts only on the north, west, and east directions for the α tubulin monomer, and only on the south, west, and east directions for the β tubulin monomer. The remaining directions correspond to the intra-dimer interface in the dimer. The fraction of native contacts (Q_n_) for a dimer is the number of contacts in a given conformation (frame number greater than 0) divided by the number of contacts in the initial structure (frame number = 0). The number of contacts for the north and south (longitudinal) interfaces are around 40-100, while the number for the east and west (lateral) interfaces is 40-80. A drop in Q_n_ to 0 indicates a break in the corresponding lateral or longitudinal interface in the lattice.

In our simulations with single point pulling, we observed two breaking events based on the drop of respective Q_n_ values. After the first breaking event due to loss of intra-dimer contacts, Q_n_ for the W lateral interface decreases to 0 resulting in the first lateral break between two PFs. The loss of the E lateral interface usually corresponds to the critical breaking event. A steep decrease in the longitudinal Q_n_ signals a longitudinal break in one of the PFs from the MT lattice (Fig. S20).

#### Protofilament Bending angles

The PF bending angle is the maximum angle obtained between two vectors characterizing the mutual orientation of three dimers from the PF (as shown in Figs. S21 and S22). In our pulling simulations, we determined the bending angles for all 13 PFs up to the point of the first longitudinal break. For each set-up, we collected the angles from all the trajectories and we built histograms (with 1° bins) using the obtained values. Furthermore, we calculated cumulative distribution functions (CDFs) of the kinking angles from the collected data and compared them with the experimental CDFs using Origin software.

##### MTs with both ends fixed

Here, for each PF in the MT lattice we obtained the largest bending angles at each time step up to the point of the first longitudinal breaking. We measured the angles at each frame on all PFs using the two vectors between the center of mass of 3 different dimers (3 points) on a PF: (1) the first dimer in the PF (Fig. S21 (A, R1)); (2) each consecutive dimer in the PF, and (3) the last dimer in the PF (Fig. S21 (C, R8)). The sign of the angle is considered positive when the PF is bending outside the MT [Szatkowski et al., 2019] and we used the absolute values of the angles for our histograms. For MT8, dimers from the rings R1 and R8 were used as reference. For MT12, we used R1-R12, R2-R11 and R3-R10, and for MT16 we used R1-R16, R2-R15, R3-R14, R4-R13 and R5-R12 as reference. We note that R3-R10 in MT12 and R5-R12 in MT16 have the same length as the MT8 lattice without the effect of fixed ends. In the case of the multi-point pulling we followed the same procedure as above but we considered angles from all the PFs with pulled dimers and we used R1-R12 and R1-R16 as reference for pulling on MT12 and, respectively, on MT16.

##### MTs with the free plus end

For the simulations of the MTs with the free plus end removal of the constraints on the plus end allows the free movement of dimers which were fixed in the above set of simulations. Therefore the previous method of calculating the bending angles for the free plus end simulations would result in a distortion of angles. For the MT8 system we did not observe this effect because on the minus side we always overestimate the measured angles (because of the fixed end) and underestimate angles on the side of the plus end. For the MT12 and MT16 lattices with free plus ends, we changed somewhat the procedure to determine the bending angles. Operationally, following our previous approach [Szatkowski et al., 2019], we obtained the angles between the following 3 selected points (Fig. S22): (1) a pseudo position created by translating the coordinates of the center of mass of the fixed dimer by 50 Å towards the minus end (Fig. S19, R); (2) the center of mass of fixed dimer (Fig. S22, R1) and (3) the center of mass of the consecutive dimer on the same PF (Fig. S22, R2 → R12). We calculated the highest angles for each PF until the first longitudinal break and we multiplied the resulting angles by 2 since these angles do not cover the entire MT. Then we followed the same method used for the fixed ends case to calculate the angles for lattices with different lengths.

## Experimental Methods

Chemical reagents purchased from Sigma. Tubulin proteins were purchased from Cytoskeleton as lyophilized powder either unlabeled or rhodamine-labeled. Tubulin was hydrated into buffer PEM-100 (100 mM PIPES, pH 6.8 using KOH, 2 mM EGTA, 2 mM MgSO_4_) and incubated for 10 minutes on ice. Hydrated tubulin was clarified by centrifugation at 366,000xg at 4°C for 10 minutes. The supernatant was supplemented with 2 mM GTP and incubated at 37°C for 20 minutes to polymerize MTs. MTswere stabilized by adding Taxol at 10 *μ*M and incubated another 20 minutes at 37°C to equilibrate Taxol. GFP-labeled, human Katanin p60 protein was purified from bacterial expression using an MBP tag for affinity purification as previously described [Belonogov et al., 2019].

MT severing assays were performed in an experimental chamber made from a slide and coverglass held together by two strips of permanent double-stick tape (3M) to create a flow path. The cover glass was pre-treated to create a hydrophobic surface through incubation in 2% dimethyldichlorosilane “Repel Silane” (GE Healthcare) after initial cleaning using UV-Ozone for 10 minutes. To specifically bind MTs to the surface and block other proteins, the chamber was flowed through with anti-tubulin antibodies MAB1864 (MilliporeSigma, Burlington, MA) at 2% (w/v) in Katanin Activity Buffer (20 mM Hepes pH 7.7, 10% glycerol, 2 mM MgCl_2_) and incubated for 5 min. The remainder of the surface was blocked using the tri-block co-polymer Pluronic F127 dissolved to 5% in Katanin Activity Buffer, and incubated for 5 min. Taxol-stabilized MTs were diluted 1:100 in Katanin Activity Buffer with supplemented Taxol and flowed into the chamber to bind MTs to the antibodies on the surface. Next, the Enzymatic Mix (20 mM Hepes pH 7.7, 10% glycerol, 2 mM MgCl_2_, 2 mM ATP, 0.025 mg/mL BSA, 0.05% F-127, 20 μM Taxol, 10 mM DTT, 15 mg/mL glucose, 0.15 mg/mL catalase, 0.05 mg/mL glucose oxidase) was flowed in to remove unbound MTs and for oxygen scavenging. MTs were imaged for 2-5 minutes prior to adding the katanin enzymes.

Severing data was recorded using a Nikon Ti-E microscope outfitted with a home-built total internal reflection fluorescence laser system with both 561 nm and 488 nm lasers for imaging MTs and GFP-katanin, respectively. The objective was 60x, NA 1.49 oil-immersion objective creating an effective pixel size of 108 nm on the Andor Ixon EM-CCD camera. To capture the dynamics of MT bending and breaking, images were recorded either in the red (MT) channel or the green (GFP-katanin) channel with no delay. We found the GFP channel had better signal to noise to determining the time when the katanin bound and the bending angles. MTs were imaged for 2-5 minutes prior to adding katanin and until all MTs were destroyed. After imaging in one location, the remainder of the coverslip was inspected to ensure that the severing enzyme affected the entire sample equally, and that the loss of polymer we observed was not due to the incident light required for imaging. Additional control experiments were performed in the absence of ATP to ensure that the enzymatic activity was responsible for the bending and breaking of MTs, and not the imaging.

## Acknowledgments

This work has been supported by National Science Foundation (NSF) Grants MCB-1412183 and MCB-1817948 to R.I.D. and Grant MCB-1817926 to J.L.R.

