## Supplemental Material for "Molecular Investigations into the Unfoldase Action of Severing Enzymes on Microtubules"

### Supplemental Results

#### SI.1 Simulations of Pulling Between Protofilaments at Different Orientations

Comparison of kinking angles from the *in-vitro* severing assays with our indentation [Jiang et al., 2017, Szatkowski et al., 2019] and pulling simulations revealed that the indentation angles are better correlated to experiments than the angles resulting from the pulling simulations. One difference between the two computational set-ups is that in pulling we apply point forces on the dimer(s) from a PF, while for indentation we can apply forces either on a PF or between PFs. We set out to determine the response of an MT lattice to pulling forces applied to a position located at the interface between 2 PFs. We also probed the orientation of the applied force. Operationally, we pulled on a selected residue (R: #38660) from the H12 helix of a β tubulin monomer, which is part of the lateral interface between PF5 and PF6 (Fig. S19A). We used the same pulling speed as before (2 μm/s) and different orientations.

We first pulled the residue perpendicular to the long axis of the MT filament in the upwards direction (+Y or +90°). It is worth noting that this position is only partly exposed to the MT lumen and thus it is unlikely to be accessible for this type of pulling by a MT bound molecular machine. This set-up results in rotation/moving of the residue R from the lateral interface to the surface of MT at ~ 80 pN, followed by the loss of intra-monomer contacts due to partial unfolding of tertiary structure. The break of the lateral contacts between PF5 and PF6 occurs at 245 pN and between PF6 and PF7 at 314 pN, followed by a longitudinal break at 247 pN. The resulting PF fragment unzips towards the minus end under a critical breaking force of ~ 326 pN resulting in the loss of 3 dimers and one β monomer.

Second, we set-up simulations mimicking indentation i.e, pulling the residue downwards (-Y or -90°) into the MT lumen, which is a possible way of action of a MT-bound molecular motor. In this case, we found the rotation of the residue from the interface into the lumen, followed by the loss of lateral contacts between PF5 and PF6 at a critical breaking force of 239 pN. This leads to the inward bending of PF6 resulting in a first longitudinal interface break at 180 pN, and the unzipping of the resulting PF6 fragment towards the minus end. We stopped the simulation when the residue touched the opposite end of the MT lumen leading to bending of the PF without losing any dimers.

In order to determine the role of force directionality in the bending of the MT lattice, next we pulled on the same residue R but at an angle of 45° in both directions (up and down). Pulling up at 45° (+Y direction), led to an initial rotation of the residue from the lateral interface to the top surface under a ~ 80 pN force, followed by the loss of intra-monomer contacts similar to the pathway for pulling up at 90°. The critical breaking force leads to the loss of lateral contacts between PFs 5 and 6 and between PFs 6 and 7 at 402 pN. Then came the break of a longitudinal interface at 301 pN, finally leading to the loss of single dimer from the lattice. This type of run thus follows pathway P1 (Fig. 1A). Pulling R in the opposite direction at 45° in -Y direction results in pulling R into the lumen at ~ 138 pN. After losing lateral contacts, the lattice loses its first longitudinal interface at the critical breaking force of 251 pN leading to the final loss of a dimer. These results show that, as expected based on our indentation results, the best orientation for pulling between PFs is perpendicular down.

When we compared the bending angle distributions from these set-ups with the experimental data, we found that pulling downwards perpendicular to the long axis of the MT lattice (**Fig. S19D**), which resembles our indentation set-up, leads to better correlation with experiments than pulling residues upwards (**Fig. S19C**) and pulling at an angle of 45$^o$ (**Fig. S19E** and **S19F)**. This validates our hypothesis that during severing indentation is more important than pulling.

#### SI.2 Detailed Description of the Multi-point Pulling Runs

#### Pulling on 3 dimers from 12 dimers long MTs

We applied pulling forces on MT12 lattices at 3 locations chosen such as three different severing enzymes could fit close together on the lattice (**Fig.S11C**) . We first simulated filaments with the ends fixed. We pulled on three dimers from three different PFs: PF 6 (D110), PF 7 (D72) and PF 8 (D34) with a speed of 2.0 μm/s. The breaking of the lattice follows pathway MP-2 (**Fig. 5**). The first event (**Fig. S12**), the unfolding of the C-terminal ends of the dimers closest to the fixed ends, happened at a force of 161 pN per pulled position. Initially, the β C-terminal end of the dimer closest to the minus end (D34) unfolds up to position 374 (i.e., H11, H11', H12 helices and the E-10 β strand are unfolded). At the same time, the α- C-terminal end of the dimer closest to the plus end (D110) unravels up to position 384 (i.e., H11, H11', and H12 helices unfold). Then PF 6 loses its lateral contacts with PF 5 leading to the formation of a crack next to the D110 pulled dimer (**Fig. S12B**), which propagates to both the ends of the lattice, and there is further unraveling of the C-terminal ends of two dimers (D34 and D72) (**Fig. S12C**). Next, we found the loss of lateral contacts between PFs 9 and 10 forming a crack that propagates from the minus to the plus end of the filament. The first longitudinal breaking event corresponds to the loss of the northern longitudinal interface of the pulled dimer on PF7 (D72) leading to its removal from the lattice (**Fig. S12D**). The break of the longitudinal interfaces at the fixed plus end of PFs 7, 8 and 9 corresponds to the critical break event at a critical breaking force of 370 pN. Next, D110 loses its longitudinal interface towards the plus end resulting in the unzipping of PF 6 towards the minus end (**Fig. S12E**). The final event corresponds to the loss of 2 dimers from PF 8 (**Fig. S12F**). The bent fragment from PF 6 seen in **Fig. S12F** will likely detach from the MT lattice, but the time for this event exceeds the 50 ms (or 1200 frames) run time in our simulation.

We also examined the same MT12 filaments with the plus end free. We selected the same 3 subunits on PF 6 (D110), PF 7 (D72) and PF 8 (D34) for pulling. In this set of simulations, the first event corresponds to the β C-terminal unfolding of D34 on PF 8 near the fixed (minus) end at a 140 pN force, followed by the unfolding of the C-terminal ends of all the pulled dimers (**Fig. S13B**). Similar to the fixed-end case, the lattice loses the lateral interface between PFs 5 and 6 leading to the formation of a crack, which propagates all the way to the fixed end (**Fig. S13C** and **S13D**). The next event consists of the break of the longitudinal interface at the plus end of D72 and the removal of this dimer from the lattice after minimal unzipping of the PF7 fragment towards the minus end. Following the loss of the lateral interface between PFs 6 and 7, the longitudinal interface at the minus end of D110 on PF 6 breaks leading to the unzipping of the resulting PF 7 fragment towards the plus end (**Fig. S13E**). The critical breaking force of 387 pN corresponds to the loss of lateral contacts between PFs 8 and 9, and the loss of 4 and, respectively, 2 dimers from the peeled off fragments of PF 6 and PF 8 (**Fig. S13F**).

#### Pulling on 3 dimers from 16 dimers long MT

We pulled on 3 dimers from 3 different PFs (**Fig. S11 D)**: PF 6 (D45), PF 7 (D98), and PF 8 (D151). The breaking pathway is MP-2. The first breaking force of 113 pN corresponded to the unfolding of helix H12 from the β C-terminal of D151, followed by the simultaneous unfolding of the C-terminal ends of D45 and D151 (the 2 dimers closest to the fixed ends). The first lateral interface break occurred between PFs 5 and 6 (**Fig. S14B**) resulting in the formation of a crack that propagated towards the plus end, which is accompanied by the unfolding of the C-terminal ends of D98 (**Fig. S14C**). Next, the loss of the lateral interface between PFs 9 and 10 led to the formation of a crack that propagated towards the minus end and to the loss of D98 from PF7 at a critical breaking force of 345 pN (**Fig. S14D** and **S14E**). The final steps involved the loss of the lateral interface between PFs 6 and 7, a longitudinal interface break enabling the unzipping of PF 6 towards the plus end, and the loss of pulled dimers from the lattice (**Fig. S14F**).

#### Pulling on 4 dimers from 3PFs of the 16 dimers long MT

First, we pulled on 4 dimers from 3 different PFs (**Fig. S11E**): PF 6 (D123), PF 7 (D85 and D163), and PF 8 (D47) thereby mimicking the addition of another severing motor on the MT lattice such that 2 motors act on subunits from the same PF. This set-up followed pathway MP-3. The initial breaking event at a 197 pN force corresponds to the unfolding of C-terminal ends in the dimers closest to the fixed ends of the filament: of both C-terminal ends of D123 and of only the β C-terminal end of D47 (**Fig. 6B**). This is followed by the unraveling of the C-terminal ends of all the pulled dimers. The next events correspond to the break of lateral interfaces: between PFs 5 and 6 (**Fig. 6C**) and between PFs 9 and 10 (**Fig. 6D**), leading to the formation of cracks that propagate towards the ends of the filament. Further increase in force leads to multiple longitudinal breaks in PF 7 resulting in the loss of two of the pulled dimers (D85 and D163) from the lattice (**Fig. 6E**). At the critical breaking force of 356 pN, the pulled PFs break from their longitudinal interfaces formed with the fixed dimers at the minus end of the lattice (**Fig. 6F**) and unzip towards the plus end.

To evaluate the role played by the correlated response of PFs in the bending and breaking of MT lattices under the action of pulling forces, in this set of simulations we kept the concentration of the modeled enzymes intact but, instead of pulling on 4 dimers located on 3 consecutive PFs, we pulled on dimers from non-neighboring PFs (**Fig. S11F)**, i.e., from PFs 6, 7 and 9 (D48, D85, D123 and D163). The breaking this time followed pathway MP-1. The first breaking event at 114 pN is due to the C-terminal unfolding of dimers on PF 6 (D85) and PF 9 (D163) (**Fig. S15D**). Then D48 on PF 9 loses its lateral contacts with the adjacent dimer located on PF 10. The critical breaking force at 206 pN corresponds to the unraveling of dimers on PFs 6 and 9, the loss of lateral contacts between D48 and D49, and the partial unfolding of the β C-terminal end of D85 from PF 7 (**Fig. S15E**). Then D48 loses one of its longitudinal interfaces leading to a minimal unzipping of PF 6 and the removal of D48 from the lattice (**Fig. S15F**).

#### Pulling on 4 dimers from 4PFs of 16 dimers long MT filaments

We pulled on 4 dimers (D47, D85, D123 and D165) from 4 consecutive protofilaments (**Fig. S11 G)**: PFs 6 to 9. The breaking pattern follows pathway MP-3. Unfolding of the

C-terminal ends of the pulled dimers on PFs 6, 8 and, respectively, 9 resulted in the first breaking event at 200 pN (**Fig. S16B**). Next, D165 from PF 9 lost its lateral contacts with the adjacent dimer on PF 10 and one of its longitudinal interfaces resulting in the detachment of this dimer from the lattice (**Fig. S16C**). We also observed the breakage of the lateral contacts between PFs 5 and 6 and between PFs 9 and 10 leading to the formation of cracks that propagated to the ends of the MT filament and the removal of the D85 dimer from the lattice (**Fig. S16D**). The critical breaking force of 400 pN led to further unraveling of the C-terminal ends of D47 and D123 (**Fig. S16E**), to the break of the longitudinal interfaces of all 4 pulled PFs from the fixed minus end and the unzipping of the resulting set of PF fragments towards the plus end (**Fig. S16F**).

#### SI.3 MT Lattice Bending

Analysis of multipoint pulling simulations for MT12 and MT16 lattices with fixed ends showed that pulling on multiple PFs led to permanent bending in other PFs (Fig. S18B). To quantify this effect, we used the initial distance between the center of mass of each dimer (i) and the center of mass of the next longitudinal dimer in the same PF, i.e., the (i+13) dimer of 85 Å, and we calculated the average distance between the centers of mass of the i and i+13 dimers along each PF during the course of simulation (Fig. S18A). The extension of each PF was obtained by the summation of all the distances between the center of mass of the dimers in the respective PF minus the original distances. We found that the two PFs (5 and 10) next to the pulled PFs (6, 7, 8 and 9) exhibit the highest degree of bending (Fig. S18B), corresponding to an average elongation per dimer of ~ 2 Å. This elongation is similar to the one observed in our previous study of MT lattices with a free plus end [Szatkowski et al., 2019]. This increase in the contour length of PFs makes them too long to fit within the fixed ends of the MT lattice resulting in their permanent bending.

**Table T1.** Results for Pulling Simulations on the various set-ups and models of MT lattices (D = dimer), number of trajectories for each model and average forces (in piconewtons) for various events occurred during pulling of MT lattice. Force required to unfold the respective C-terminal ends and average maximum forces for the simulations are provided in C-terminal unfolding and critical breaking column. The force required to break the first longitudinal interface and final longitudinal break (LONG) resulting in loss of subunits are given last two columns.

| Set-up | Model | # trajectori -es | Forces (pN) | | | |
|---|---|---|---|---|---|---|
| | | | C-terminal Unfolding | Critical Breaking | First Breaking (LONG) | Second Breaking (LONG) |
| Fixed Ends | 8D, Away, Beta | 10 | 149 | 353 | 302 | 275 |
| | 8D, Away, Both | 11 | 137 | 325 | 284 | 215 |
| | 8D, Seam, Beta | 11 | 162 | 345 | 296 | 207 |
| | 8D, Seam, Both | 10 | 180 | 421 | 340 | 215 |
| | 12D, Away, Beta | 10 | 168 | 364 | 324 | 267 |
| | 12D, Away, Both | 10 | 249 | 433 | 387 | 263 |
| | 12D, Seam, Beta | 10 | 197 | 350 | 309 | 238 |
| | 12D, Seam, Both | 10 | 224 | 394 | 347 | 232 |
| | 16D, Away, Beta | 2 | 172 | 360 | 322 | 250 |
| | 16D, Away, Both | 3 | 268 | 426 | 377 | 230 |
| Free Plus End | 8D, Away, Beta | 5 | 178 | 363 | 297 | 194 |
| | 8D, Away, Both | 5 | 214 | 375 | 291 | 129 |
| | 8D, Seam, Beta | 7 | 113 | 404 | 375 | 287 |
| | 8D, Seam, Both | 5 | 315 | 510 | 442 | 294 |
| | 8D, missing 2 longitudinal dimers, pulling on D44, Beta | 5 | 147 | 496 | 392 | 286 |

| | | | | | | |
|---|---|---|---|---|---|---|
| **Defects** | 8D, missing 2 longitudinal dimers, pulling on D44, Both | 5 | 176 | 518 | 439 | 236 |
| | 8D, missing 2 longitudinal dimers, pulling on D57, Beta | 5 | 155 | 254 | 249 | 139 |
| | 8D, missing 2 longitudinal dimers, pulling on D57, Both | 5 | 341 | 341 | 304 | 169 |
| | 8D, missing 2 lateral dimers, pulling on D57, Beta | 6 | 136 | 469 | 441 | 285 |
| | 8D, missing 2 lateral dimers, pulling on D57, Both | 5 | 292 | 501 | 424 | 209 |

**Table T2.** Results for multipoint pulling simulations on the various set-ups and models of MT lattices (D = dimer), number of trajectories for each model and average forces (in piconewtons) for various events occurred during pulling of MT lattice. Force required to unfold the respective C-terminal ends and average maximum forces for the simulations are provided in C-terminal unfolding and critical breaking column.

| Set-up | Model | # trajectories | Forces (pN) | |
|---|---|---|---|---|
| | | | C-terminal Unfolding | Critical Breaking |
| Multi point pulling | 12D, Away, 3Pt | 3 | 161 | 370 |
| | 12D, Away, Free plus end, 3Pt | 5 | 140 | 387 |
| | 16D, Away, 3Pt | 1 | 113 | 358 |
| | 16D, Away, 4Pt (PFs 6778) | 3 | 197 | 356 |
| | 16D, Away, 4Pt (PFs 6779) | 1 | 114 | 202 |
| | 16D, Away, 4Pt (PFs 6789) | 3 | 200 | 399 |

#### Supplemental Figures

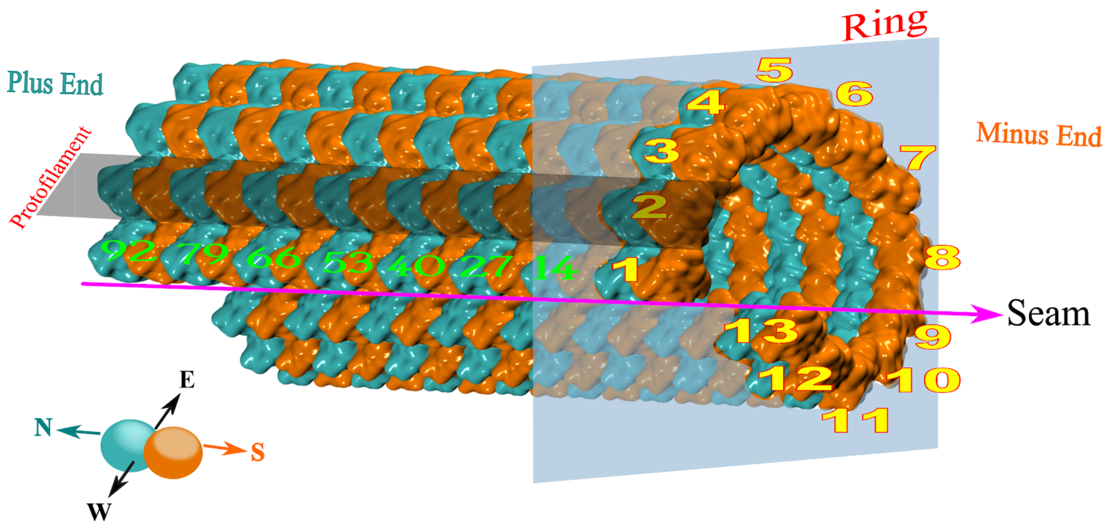


**Fig. S1:** Structure of an 8 dimers long MT lattice with 104 subunits. The MT lattice consists of 8 rings with 13 dimers (yellow) bound laterally. Dimers in a ring are sequentially numbered from i, (i+1), to (i+12), with the ends of a ring joined together at seam (magenta). The rings are joined longitudinally along the lattice from the minus end to the plus end. The dimers (green) in the first protofilament are numbered explicitly. The spatial orientation for neighbors of a given PF or dimer in the lattice (N, S, E, W) is also provided.

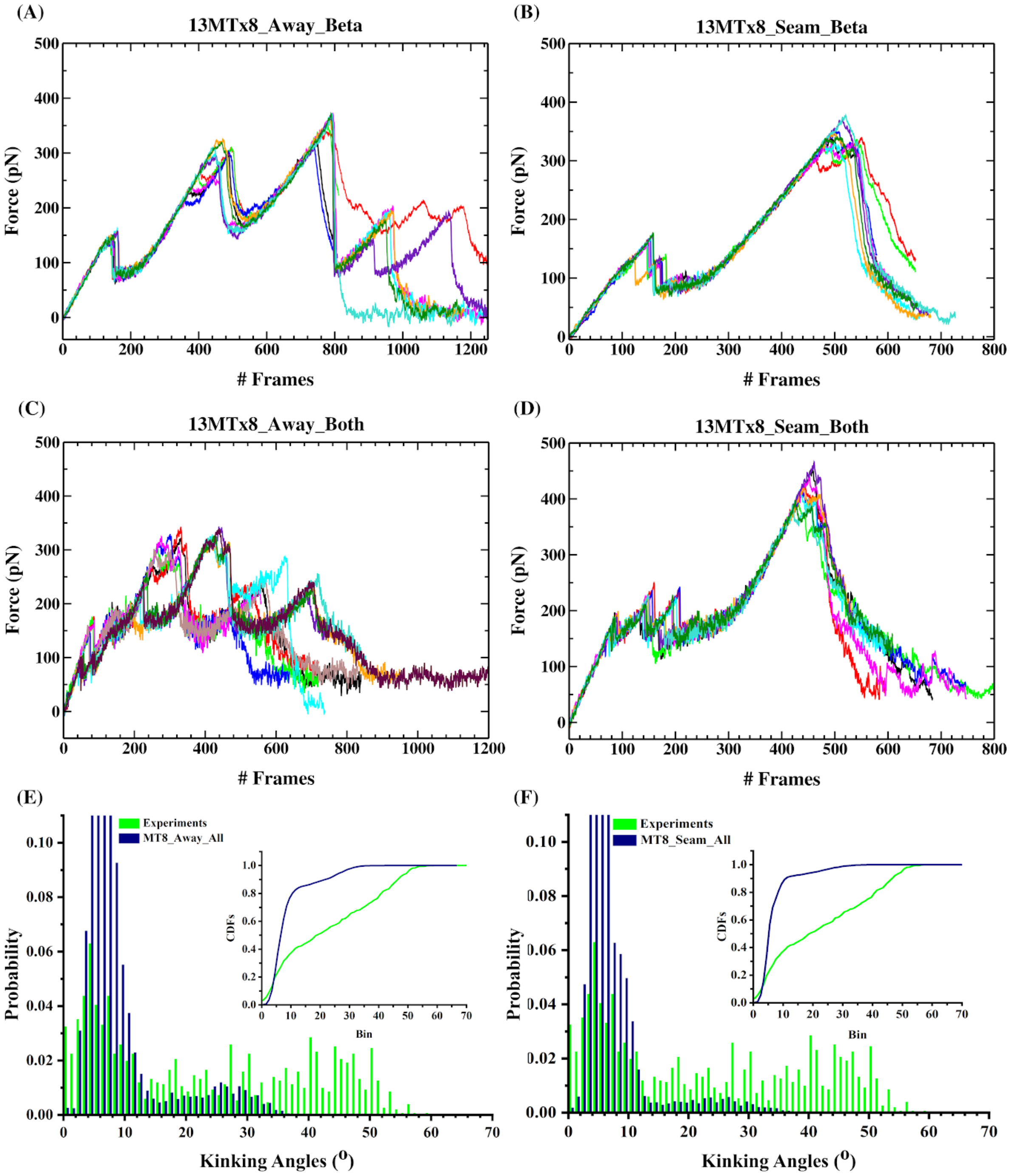


**Fig. S2:** Force distributions vs frame number (1 frame = 0.04 ms) of 8 dimers long 13 PF MT, pulling on C-terminal region of β monomer (A) away from seam & (B) on seam; α and β (C) away from seam and (D) on seam. Comparison of bending angle distributions from experimental severing assays (Green) and from simulations (Blue) for (E) away and (F) on the seam. The insets show the respective CDFs.

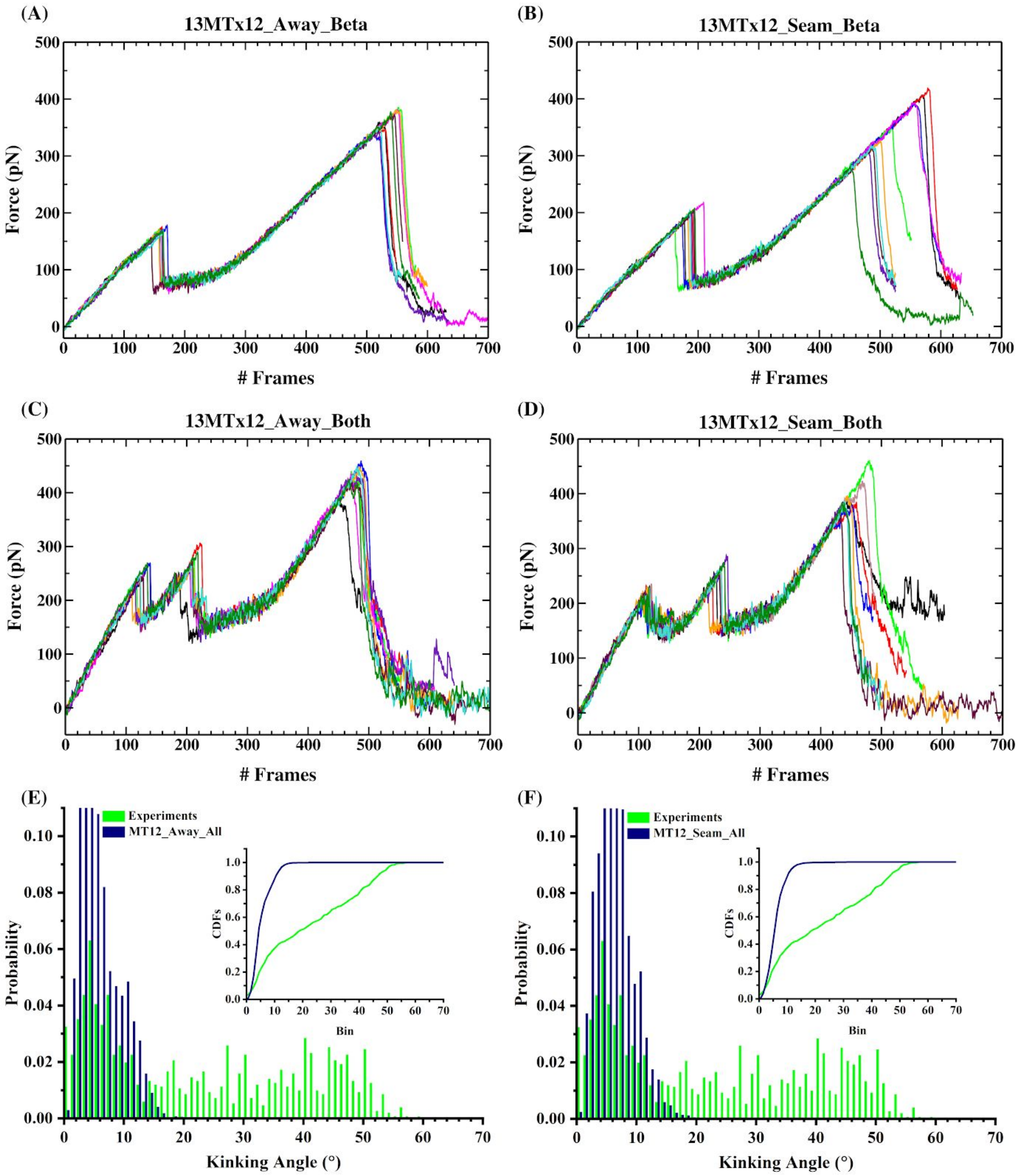


**Fig. S3:** Force distributions vs frame number (1 frame = 0.04 ms) of 12 dimers long 13 PF MT, pulling on C-terminal region of β monomer (A) away from seam and (B) on seam; α and β (C) away from seam and (D) on seam. Comparison of bending angle distributions from experimental severing assays (Green) and from simulations (Blue) for (E) away and (F) on the seam. The insets show the respective CDFs.

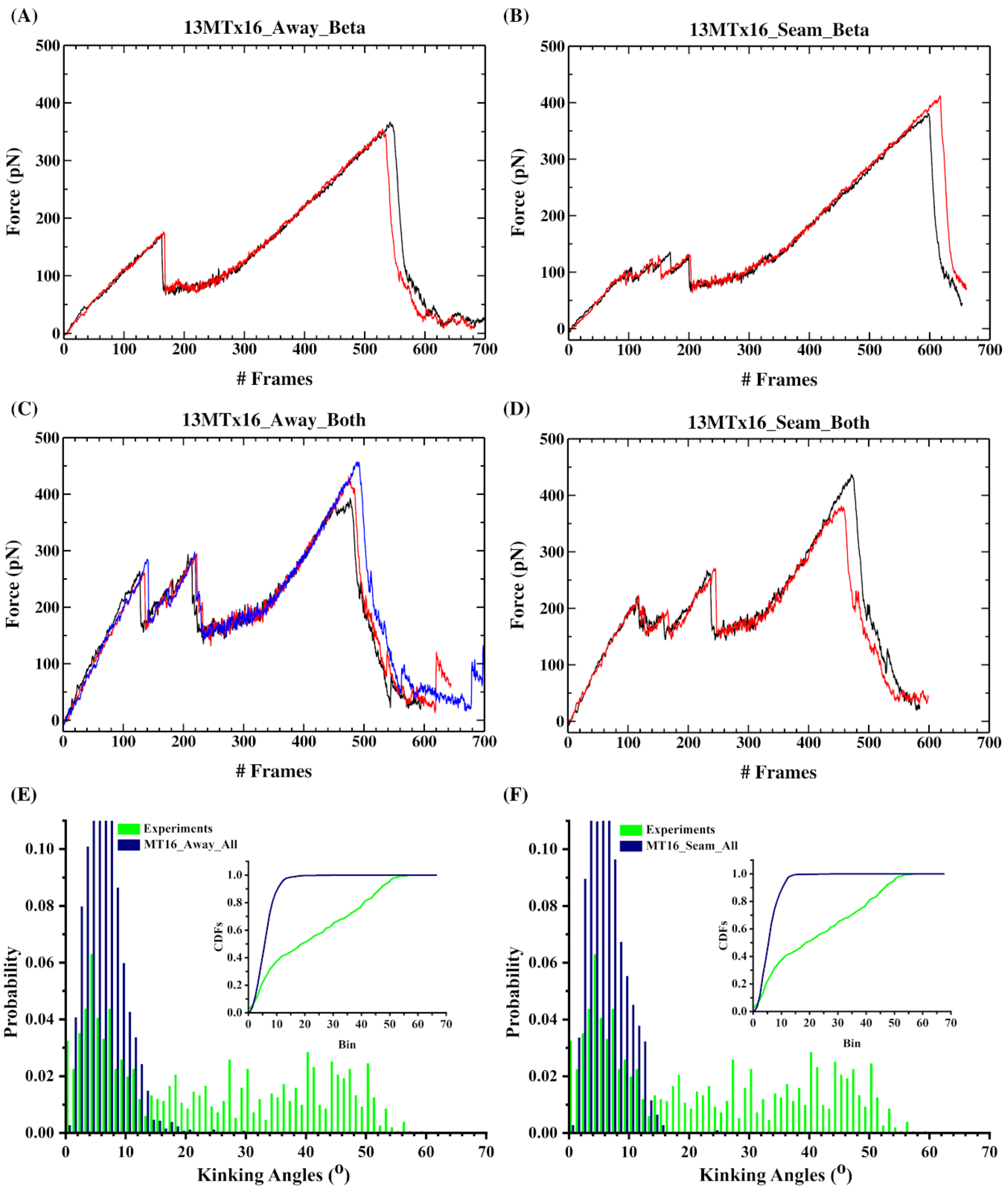


**Fig. S4:** Force distributions vs frame number (1 frame = 0.04 ms) of 16 dimers long 13 PF MT, pulling on C-terminal region of β monomer (A) away from seam and (B) on seam; α and β (C) away from seam and (D) on seam. Comparison of bending angle distributions from experimental severing assays (Green) and from simulations (Blue) for (E) away and (F) on the seam. The insets show the respective CDFs.

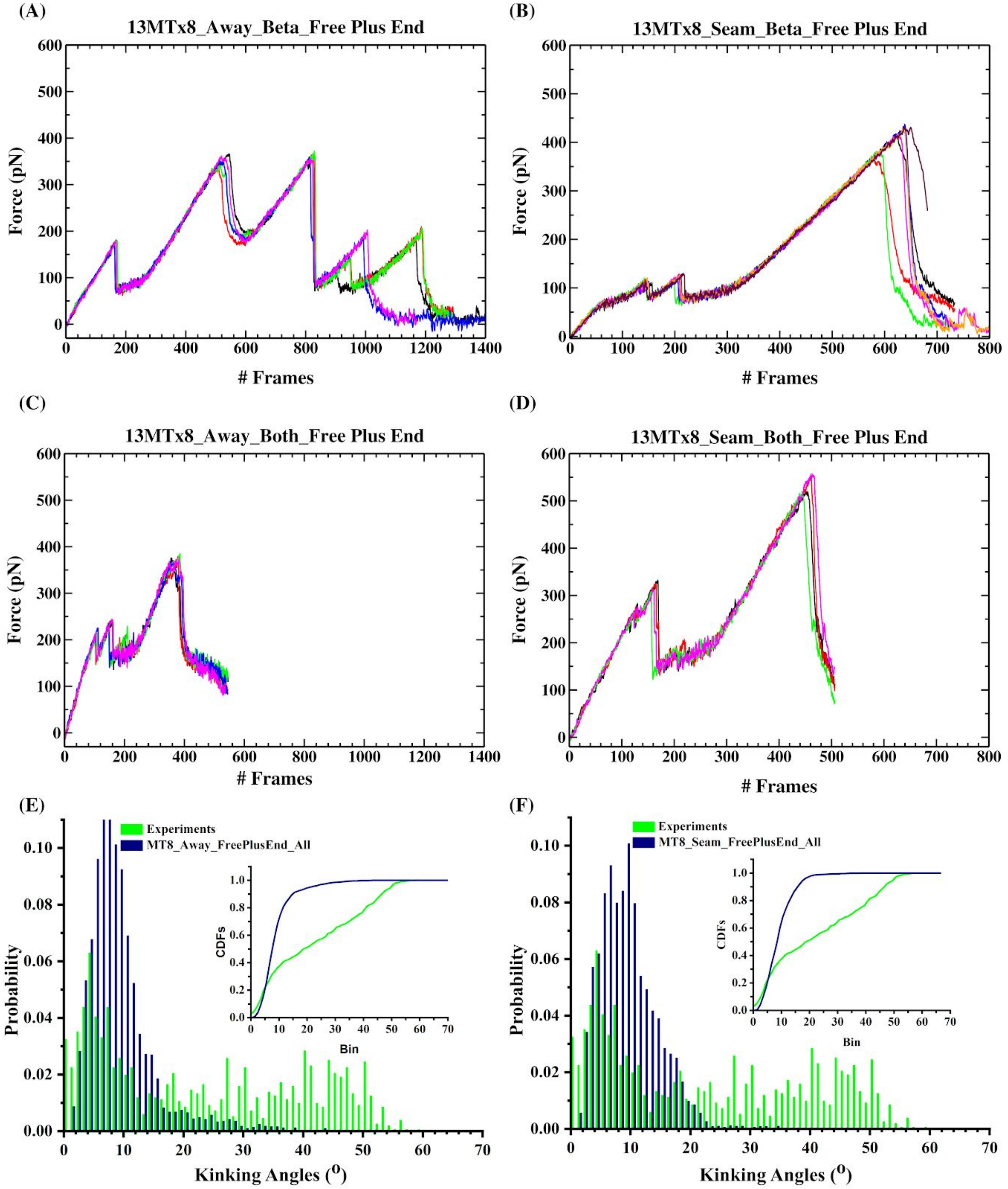


**Fig. S5:** Force distributions vs frame number (1 frame = 0.04 ms) of 8 dimers long 13 PF MT with free plus end, pulling on C-terminal region of β monomer (A) away from seam and (B) on seam; α and β (C) away from seam and (D) on seam. Comparison of bending angle distributions from experimental severing assays (Green) and from simulations (Blue) for (E) away and (F) on the seam. The insets show the respective CDFs.

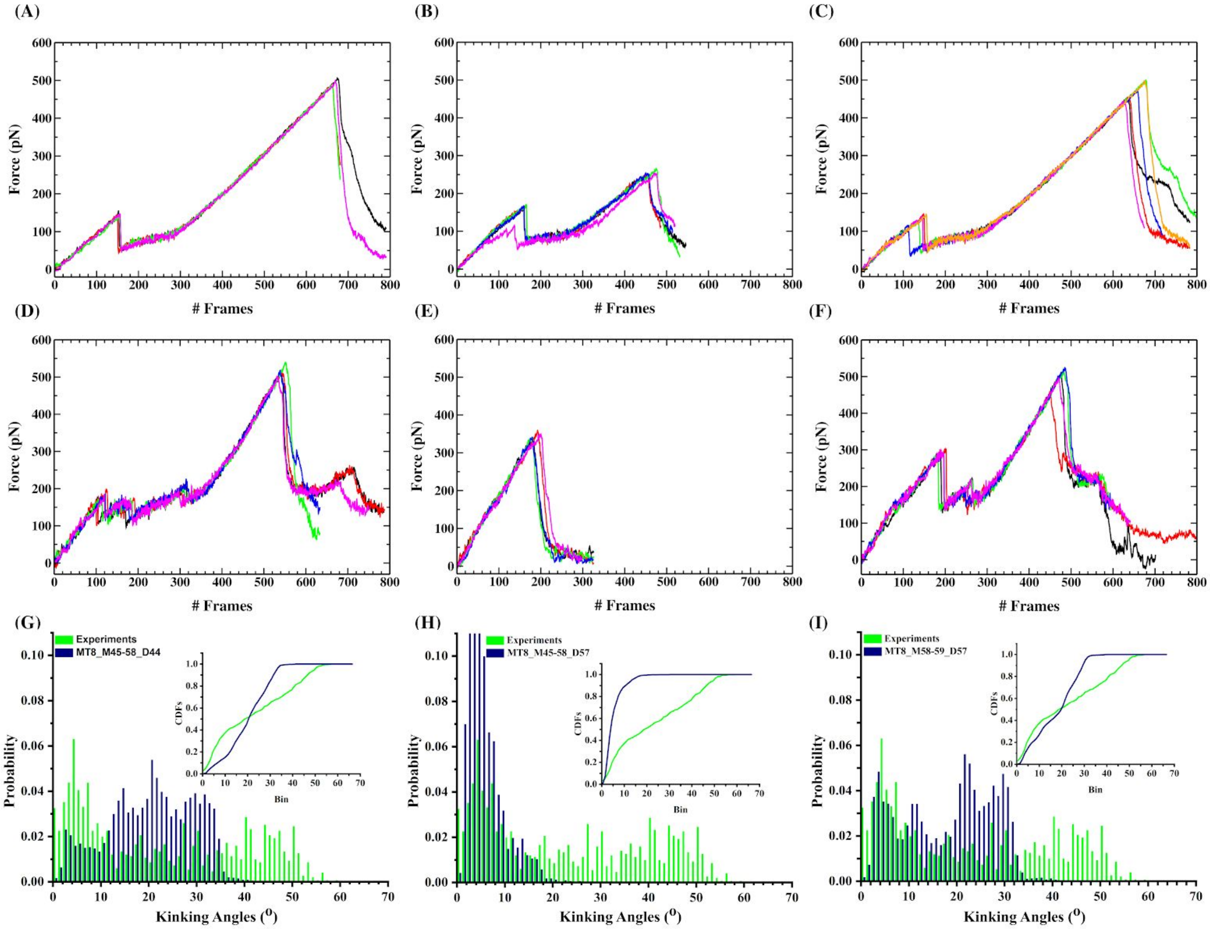


**Fig. S6:** Force evolution vs frame number (1 frame = 0.04 ms) of 8 dimers long MTs with 2% defects, with two longitudinal dimers (D45 and D58) missing and pulling on the C-terminal region of D44 (A) β and (D) α and β; D57 (B) β and (E) α and β. With two lateral dimers (D58 and D59) missing and pulling on C-terminal region of D57 (C) β and (F) α and β. Bending angle distributions from experimental severing assays (Green) and from above the simulation setups (Blue) for lattice with defects (G, H and I).

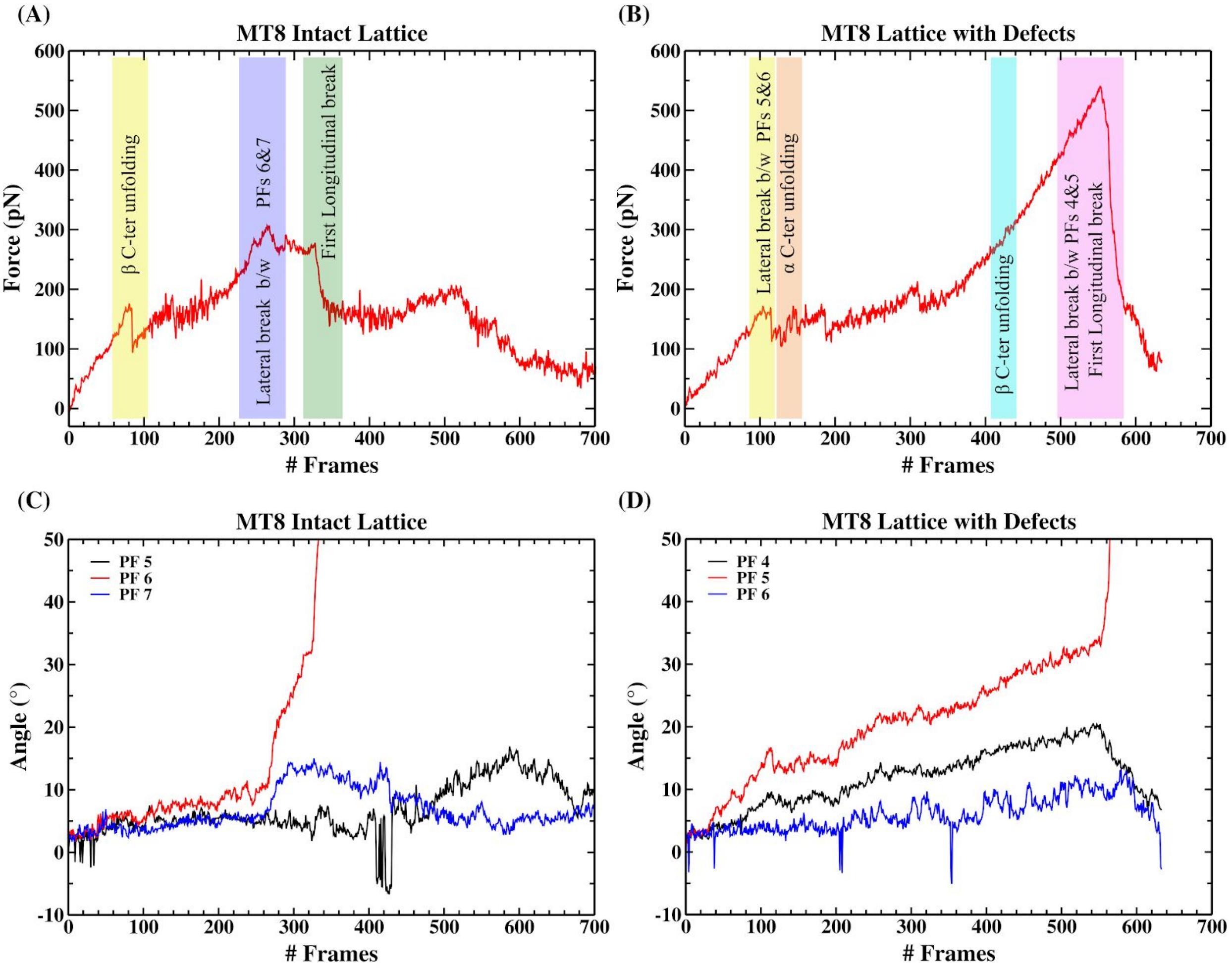


**Fig. S7** : Comparison of the time evolution of forces and of the PF bending angles along the course of pulling simulations (1 Frame = 0.04 ms) on 8D long MT (A and C) for defect-free lattices and (B and D) for lattice with defects (2 longitudinal dimers missing). Bending angles are shown for the pulled PF (PF6 in the intact lattice and PF 5 in the lattice with defects) - Red, the PF sharing the western interface with the pulled PF - Black and the PF sharing the eastern interface with the pulled PF - Blue. As described in the text, we collected the bending angles only up to the time of the first longitudinal interface breaking.

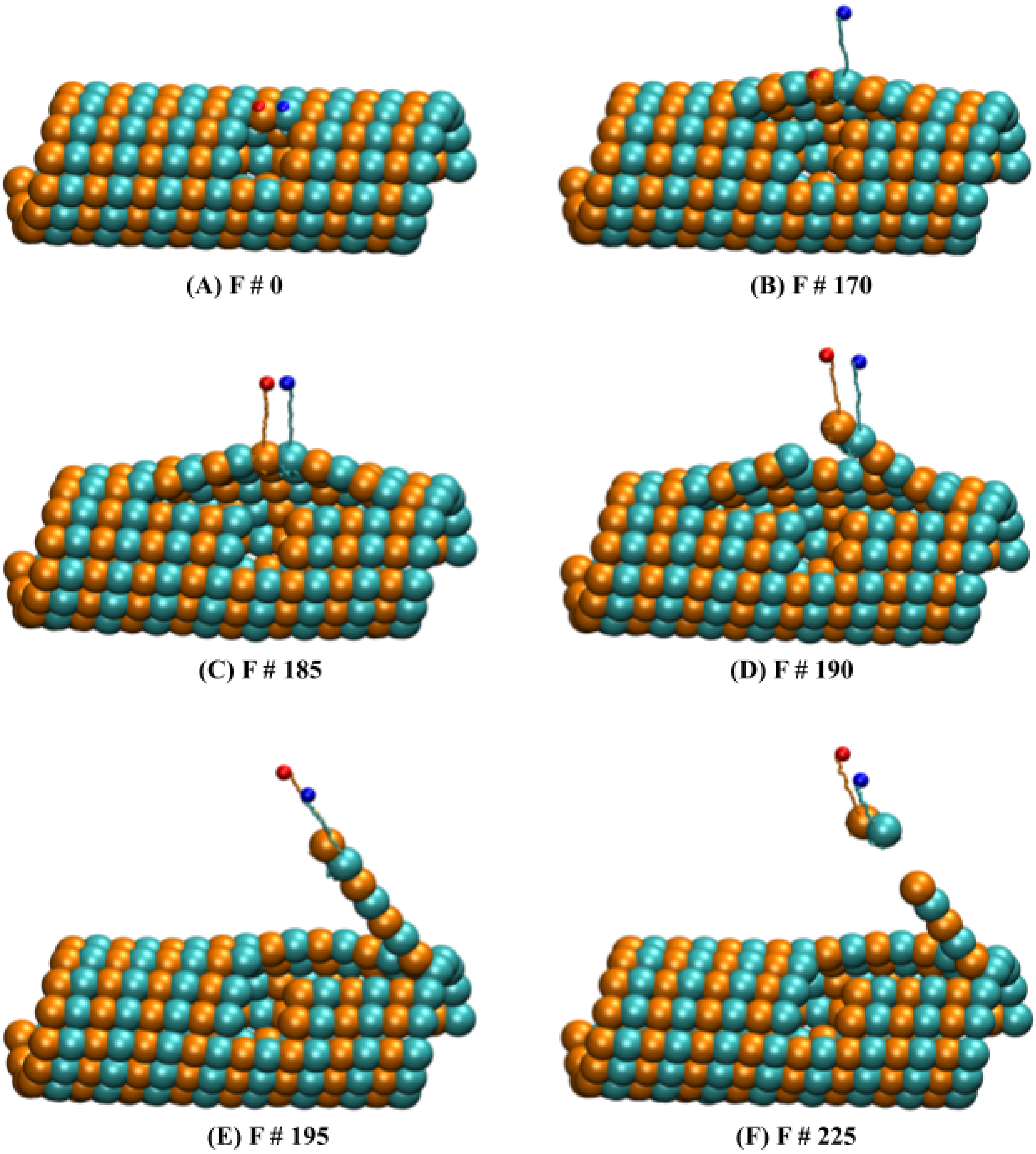


**Fig. S8:** Representative states for the MT8 lattice with two lateral dimers (D58 and D59) missing when pulled on D57. (A to F) showing different states (F # - frame number) along pathway-1 (P1), where the lattice lost a dimer upon pulling on α (Red) and β (Blue) C-terminal residue.

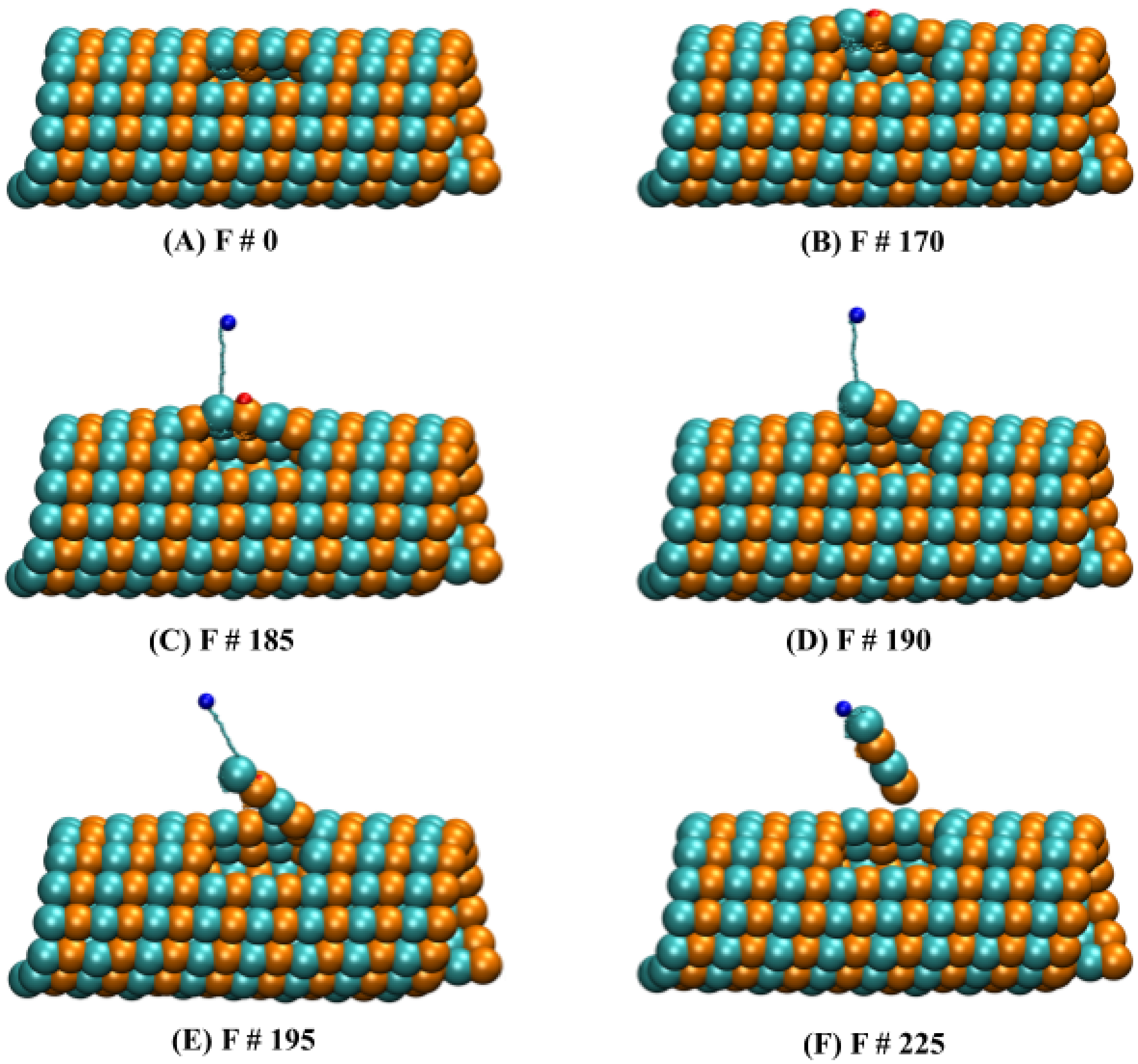


**Fig. S9:** Representative states for the MT8 lattice with two longitudinal dimers (D45 and D58) missing and pulled on D57. (A to F) showing different states (F # - frame number) along the pulling pathway-2 (P2), where the lattice lost 2 dimers upon pulling on the β C-terminal residue (Blue).

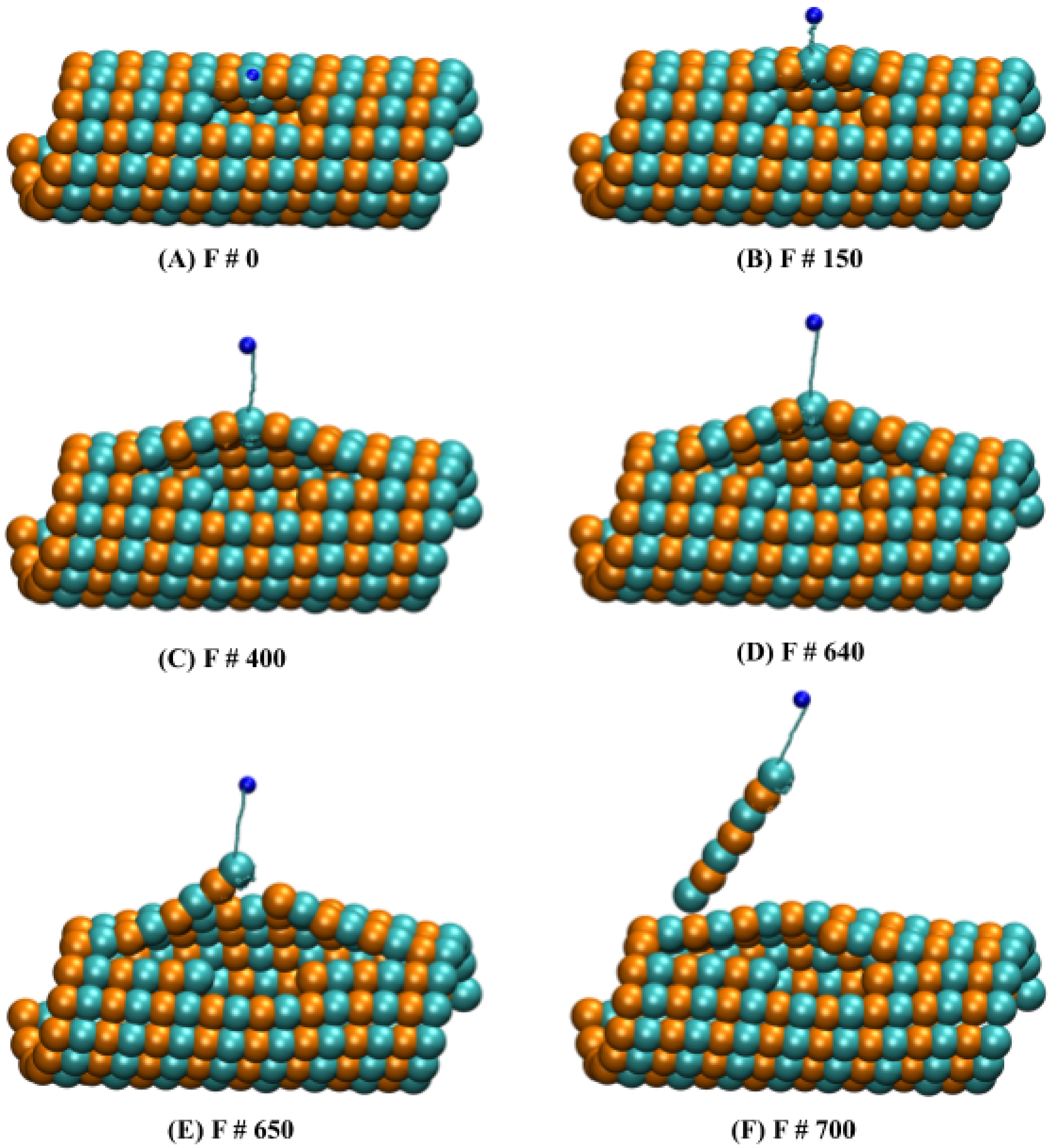


**Fig. S10:** Representative states for the MT8 lattice with two longitudinal dimers (D45 and D58) missing when pulled on D44. (A to F) showing different states (F # - frame number) along the pulling pathway-3 (P3), where the lattice lost 3 dimers upon pulling on the β C-terminal end (Blue).

### Multi point pulling

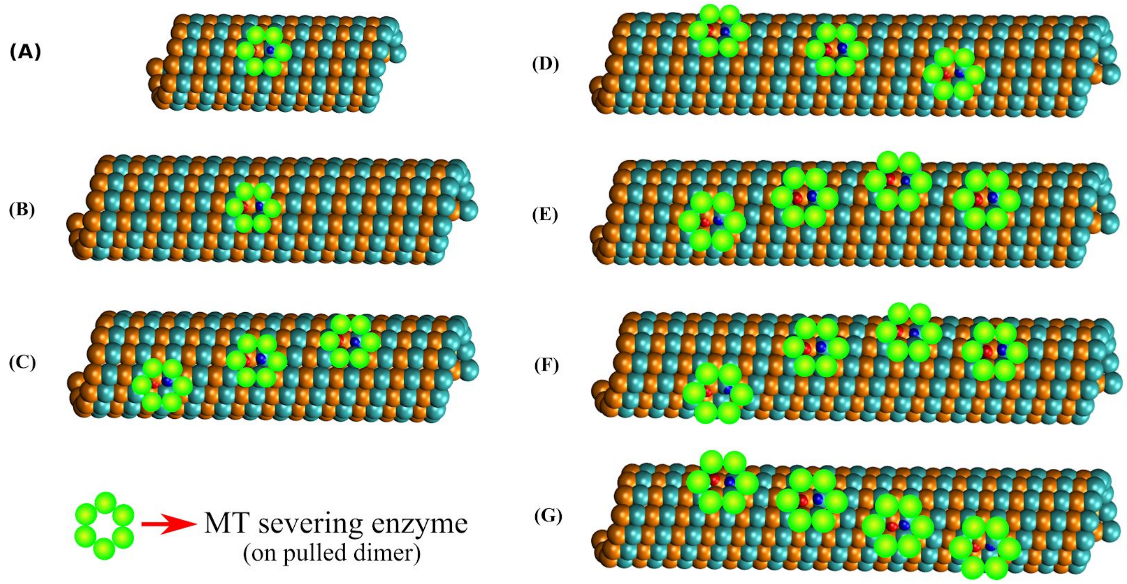


**Fig. S11:** Different set-ups for simulations mimicking the pulling action of a severing enzyme (Green) on **a** MT lattice **(A)** Single dimer pulling on the MT8 lattice (fixed and free plus end), (B) Single dimer pulling on the MT12 lattice (fixed and free plus end), (C) 3 dimer pulling on the MT12 lattice (fixed ends), (D) 3 dimer pulling on the MT16 lattice (fixed ends), (E) 4 dimer pulling on PFs 6, 7, 7 and 8 of the MT16 lattice (fixed ends), (F) 4 dimer pulling on PFs 6, 7, 7 and 9 of the MT16 lattice (fixed ends), and (G) 4 dimer pulling on PFs 6, 7, 8 and 9 of the MT16 lattice (fixed ends).

### MT12, 3 Point pulling, Fixed ends

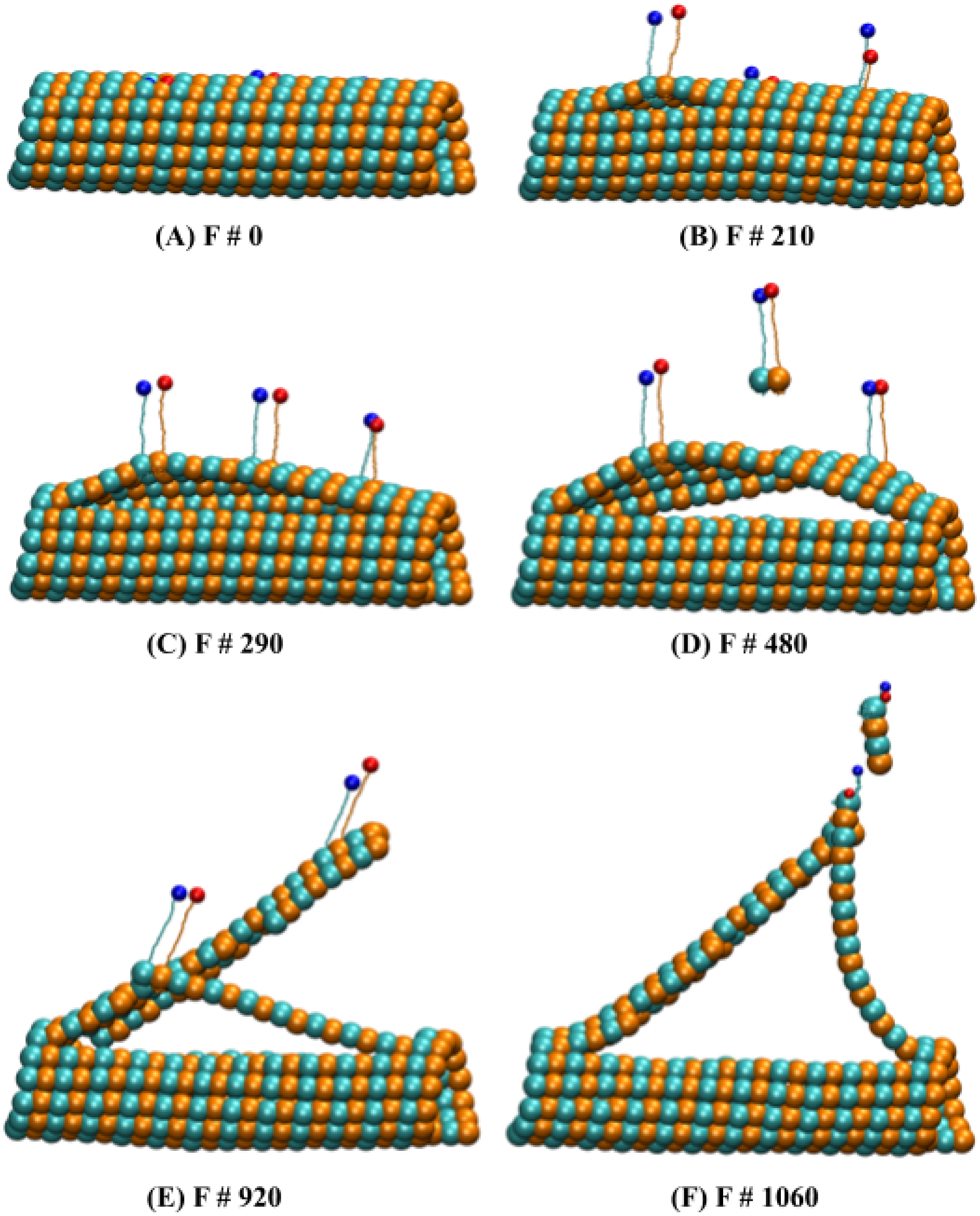


**Fig. 12:** 3 point pulling on 12 dimers long MT lattice. (A to F) showing different states (F # - frame number; 1 frame = 0.04 ms) along the trajectory. Here we pulled on the β (Blue) and α (Red) C-terminal ends of dimers D34, D72 and D110.

### MT12, 3 Point pulling, Free plus end

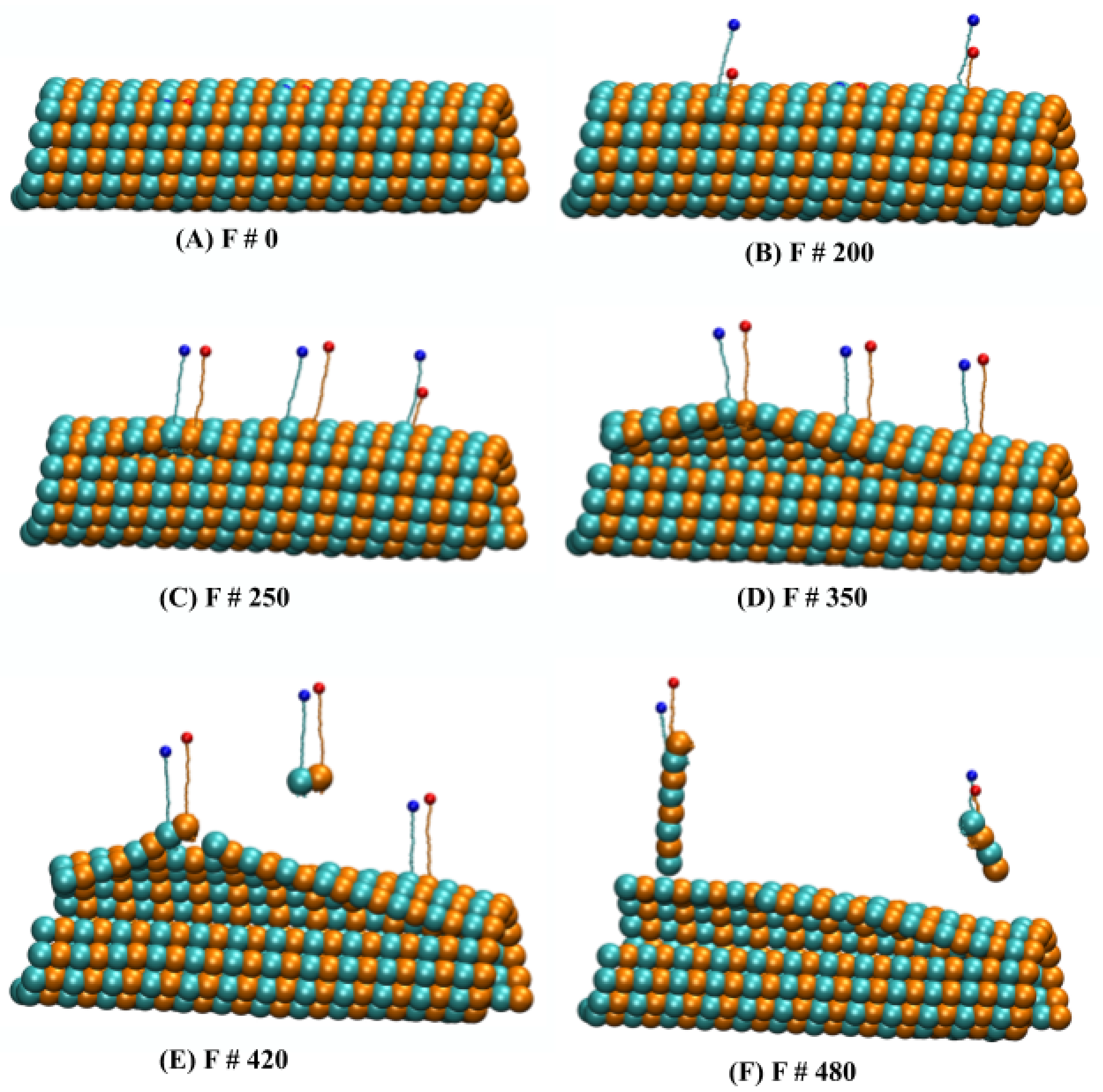


**Fig. S13:** Representative states for the 3 point pulling on the MT12 lattice with the free plus end. (A to F) showing different states (F # - frame number; 1 frame = 0.04 ms) along the trajectory. Here we pulled on the β (Blue) and α (Red) C-terminal ends of dimers D34, D72 and D110.

#### MT16, 3 Point pulling

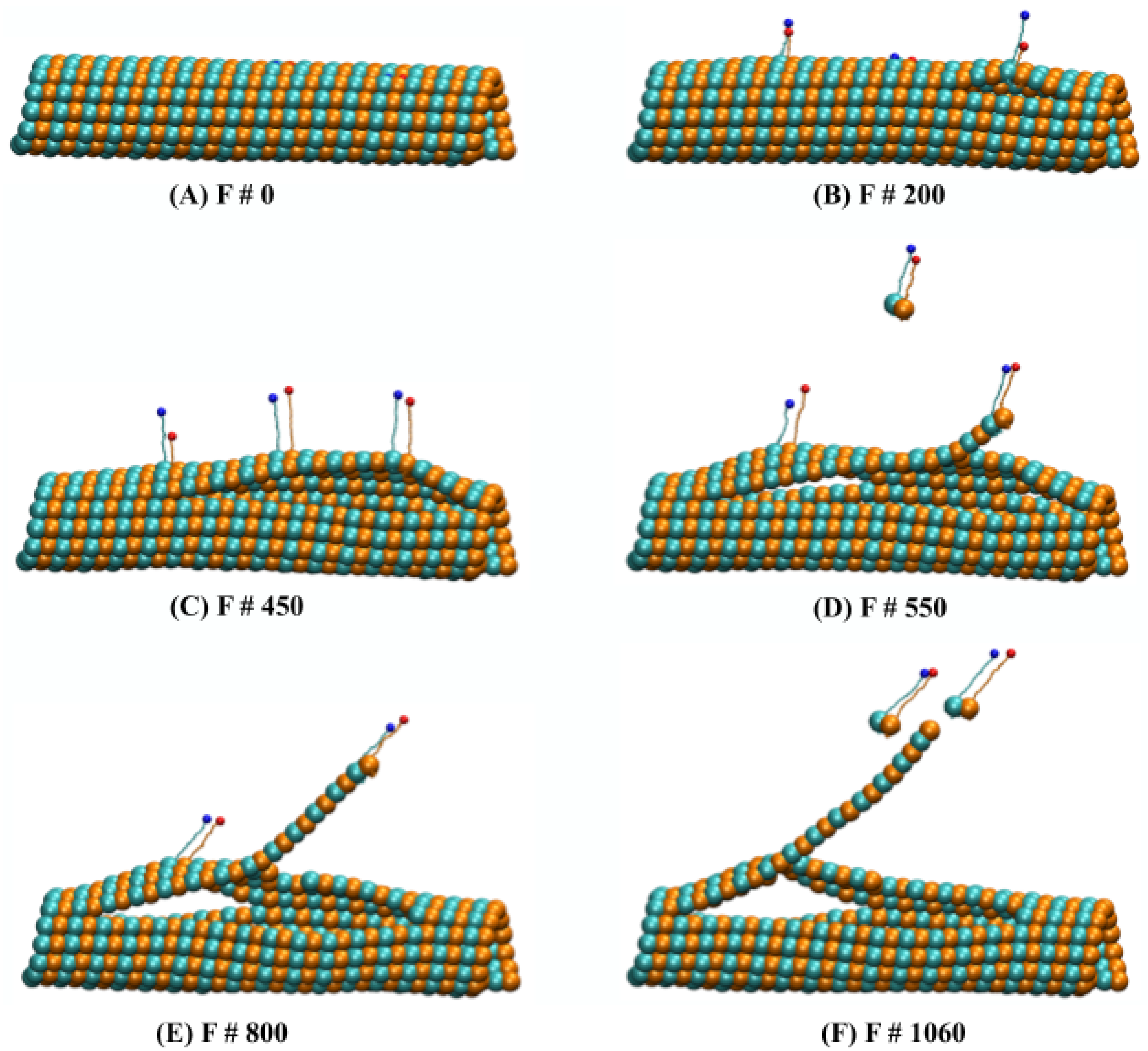


**Fig. S14:** Representative states for the 3 point pulling on PF 6, 7 and 8 of the MT16 lattice. (A to F) showing different states (F # - frame number; 1 frame = 0.04 ms) along the trajectory. Here we pulled on the β (Blue) and α (Red) C-terminal ends of dimers D45, D98 and D151.

### MT16, 4 Point pulling, PFs-6, 7, 7&9

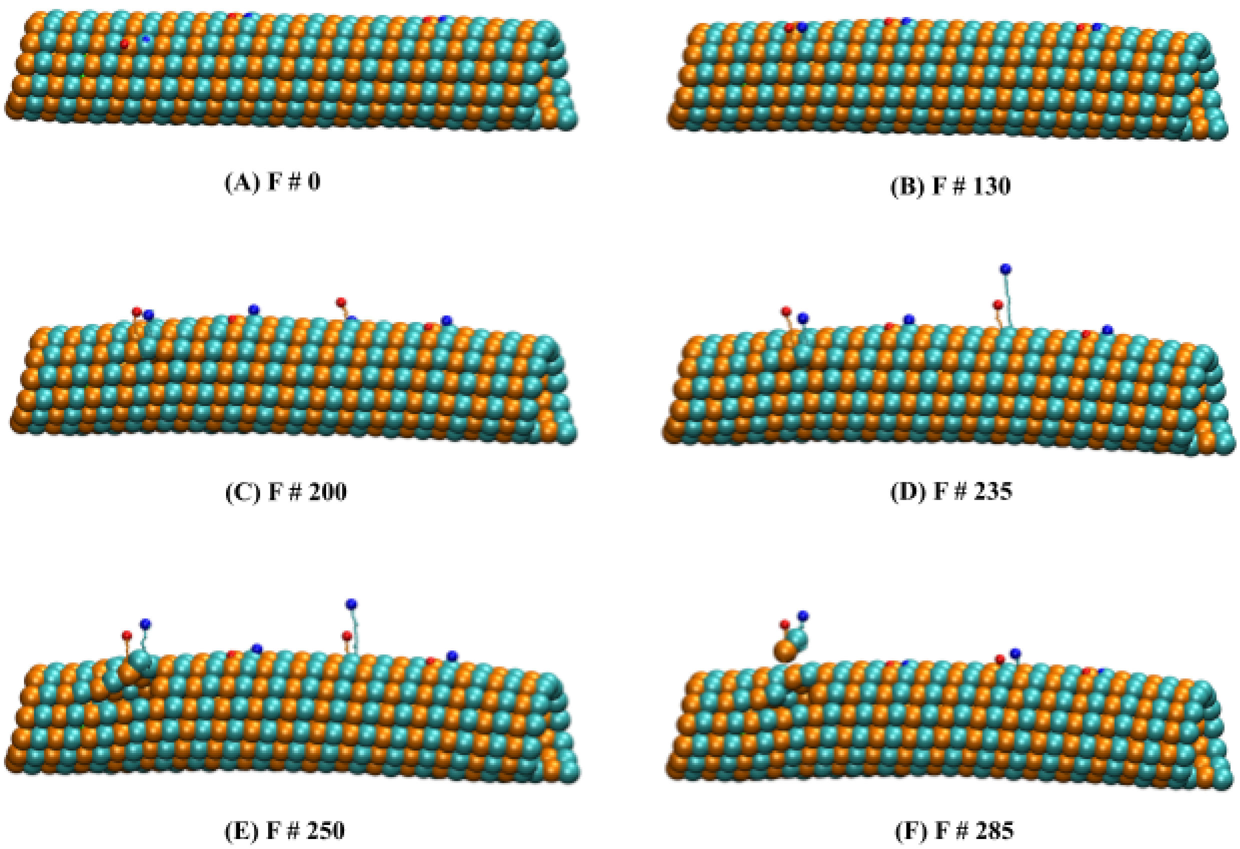


**Fig. S15:** Representative states of the 4 point pulling on PF 6, 7, 7 and 9 from the MT16 lattice. (A to F) showing different states (F # - frame number; 1 frame = 0.04 ms) along the trajectory. Here we pulled on the β (Blue) and α (Red) C-terminal ends of dimers D48, D85, D123 and D163.

### MT16, 4 Point pulling, PFs-6, 7, 8 &9

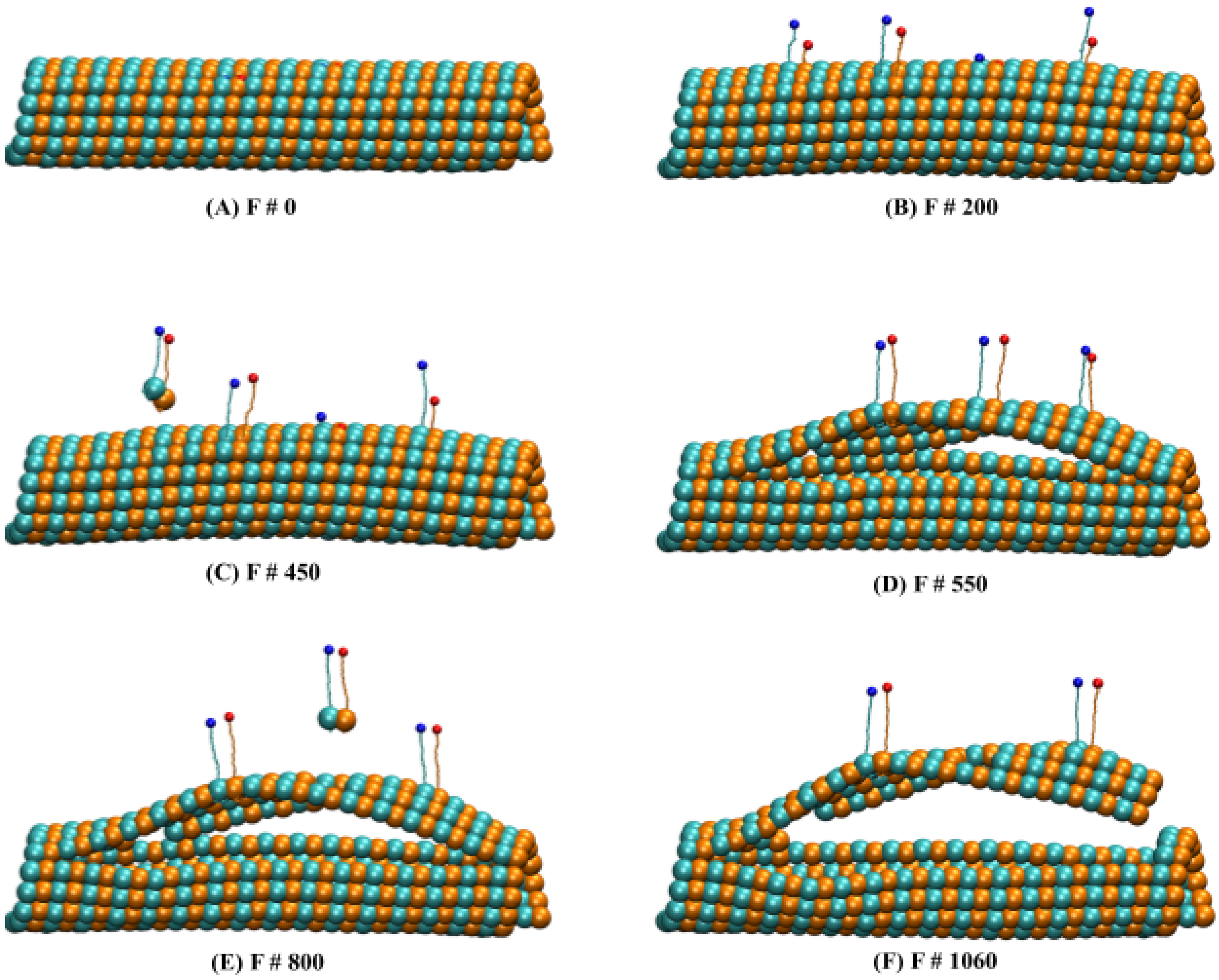


**Fig. S16:** Representative states of the 4 point pulling on PF 6, 7, 8 and 9 from the MT16 lattice. (A to F) showing different states (F # - frame number; 1 frame = 0.04 ms) along the trajectory. Here we pulled on the β (Blue) and α (Red) C-terminal ends of dimers D47, D85, D123 and D165.

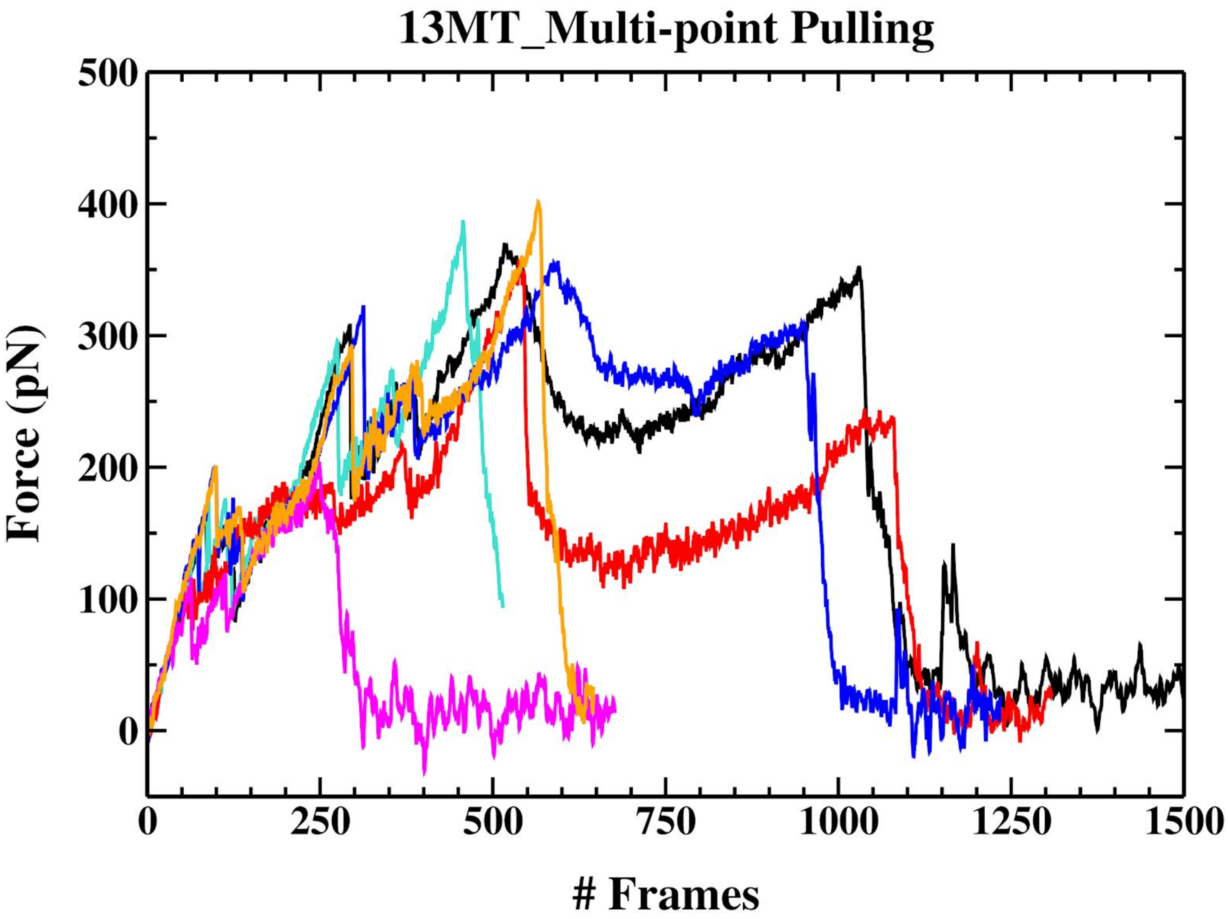


**Fig. S17:** Force distributions vs frame number (1 frame = 0.04 ms) for the MT lattices pulled on multiple dimers. (1) 3 point pulling on the 12 dimers long MT lattice with fixed ends (Black), (2) 3 point pulling on the MT12 lattice with free plus end (Red), (3) 3 point pulling on the MT16 lattice with fixed ends (Green), (4) 4 point pulling on PFs 6, 7, 7 and 8 of the MT16 lattice with fixed ends (Blue), (5) 4 point pulling on PFs 6, 7, 7 and 9 of the MT16 lattice with fixed ends (Magenta), and (6) 4 point pulling on PFs 6, 7, 8 and 9 of the MT16 lattice with fixed ends (Orange).

#### Protofilament elongation analysis

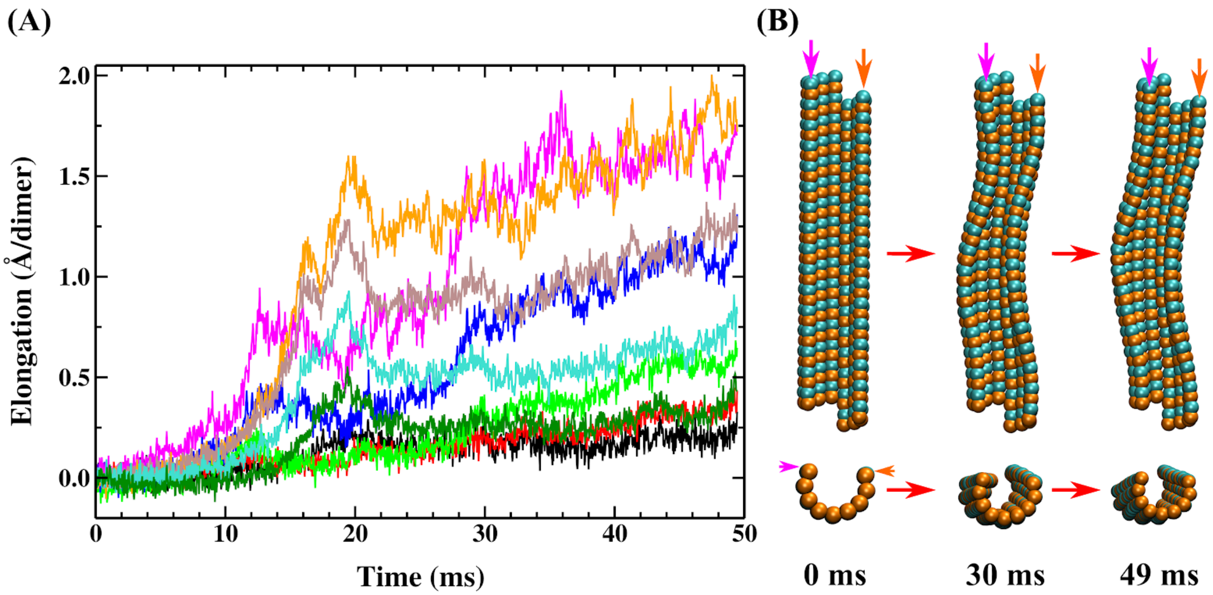


**Fig. S18: (A)** Elongation of protofilaments (PFs) for multi-point pulling of the MT16 lattice with fixed ends. Protofilaments: 1-black, 2-red, 3-green, 4-blue, 5-magenta, 10-orange, 11-brown, 12-cyan and 13-dark green. **(B)** Representation of PFs bending during simulation: PFs (5-magenta, 10-orange) next to pulled PFs.

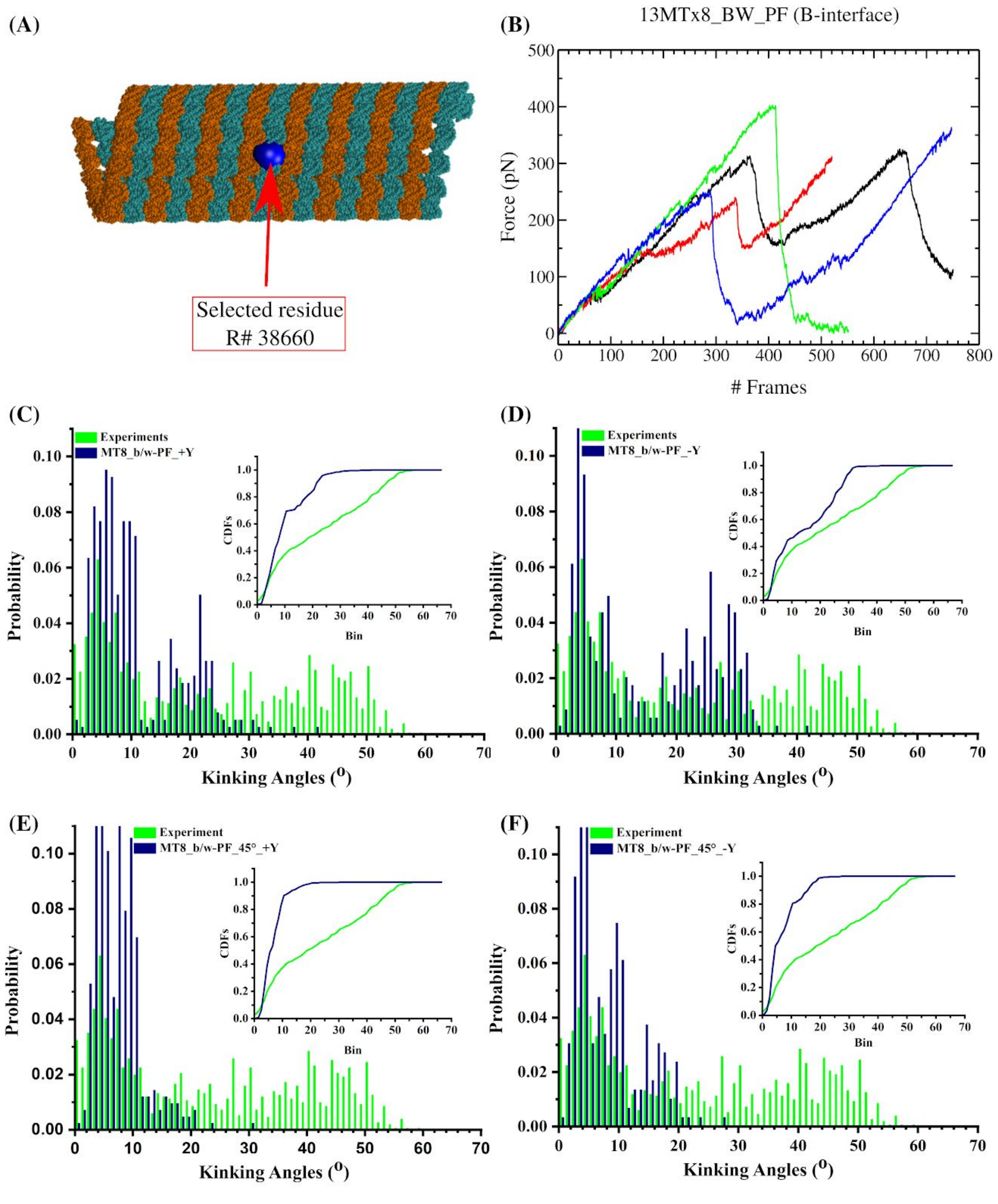


**Fig. S19:** (A) Set-up for an MT lattice pulled between PFs on a single position (Blue), (B) Force distributions vs frame number (1 frame = 0.04 ms) for the MT8 lattice pulled between PFs in the directions: perpendicular upward (+Y, $90^{o}$ , black), perpendicular downward (-Y, $-90^{o}$ , red), upwards at $45^{o}$ ($45^{o}$+Y , green) and downwards at $45^{o}$ ($45^{o}$-Y , blue). Bending angle distributions from the experimental severing assays (green) and from the simulations (blue) of pulling on residue R between PFs at (C) $90^{o}$ (+Y), (D) $-90^{o}$ (-Y), (E) $45^{o}$ (45+Y) and (F) $-45^{o}$ ($45^{o}$-Y). Insets represent the respective CDFs.

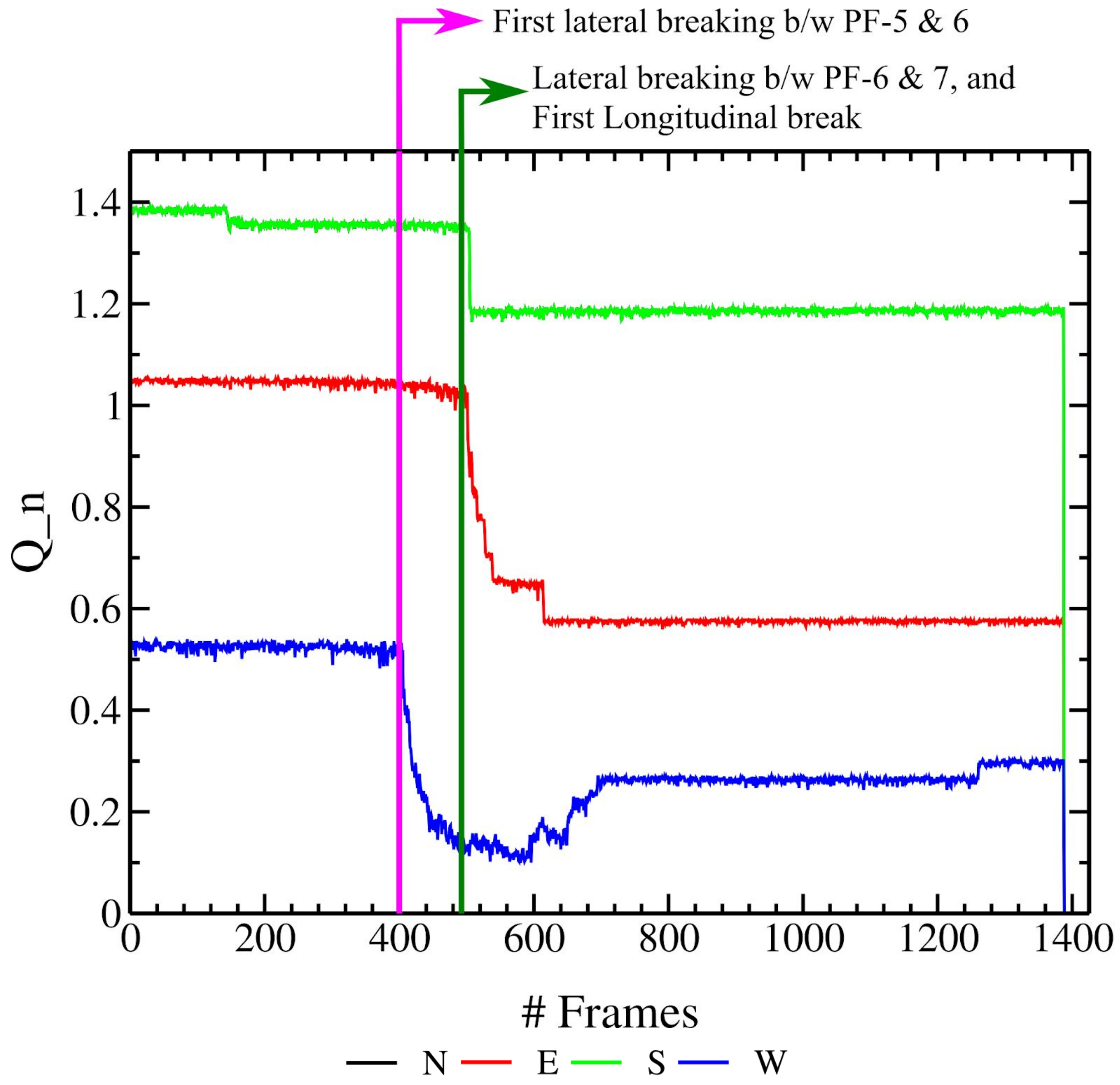


**Fig. S20:** Example of the time evolution of native contacts for a MT8 pulling simulation. Y-axis shows the $Q_n$, i.e, the total number of contacts at a particular frame (1 frame = 0.04 ms) for an interface divided by the average number of native contacts for the same interface in the starting configuration (at frame 0).

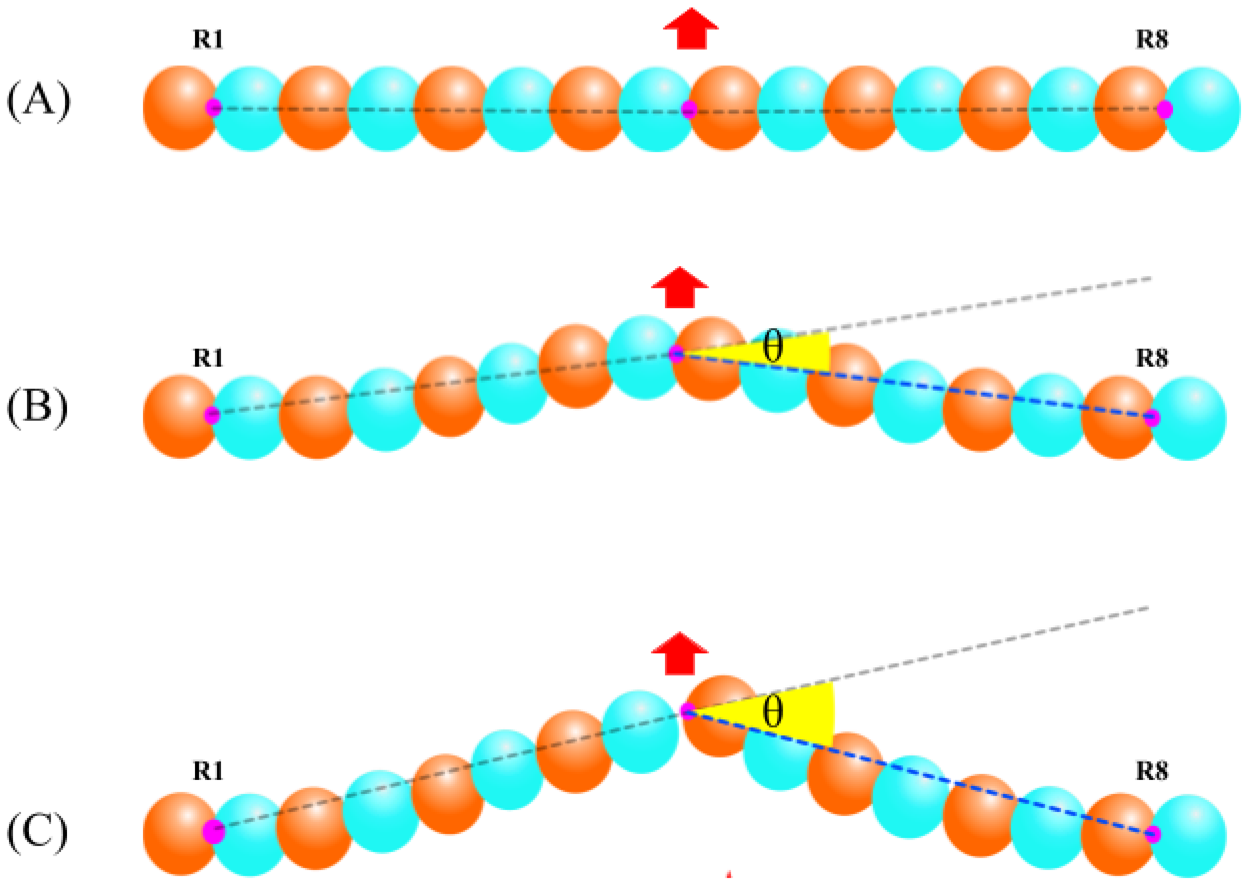


**Fig. S21:** Representation of bending angles measurements for a PF with fixed ends. Orange and cyan spheres represent α and β tubulin monomers. R1 and R8 are the fixed end dimers. Red arrow indicates the pulling point on PF and magenta indicates the center of mass of the respective dimers. (A) initial configuration (frame 0), (B) intermediate configuration as a result of pulling and (C) configuration corresponding to the first longitudinal break in the PF.

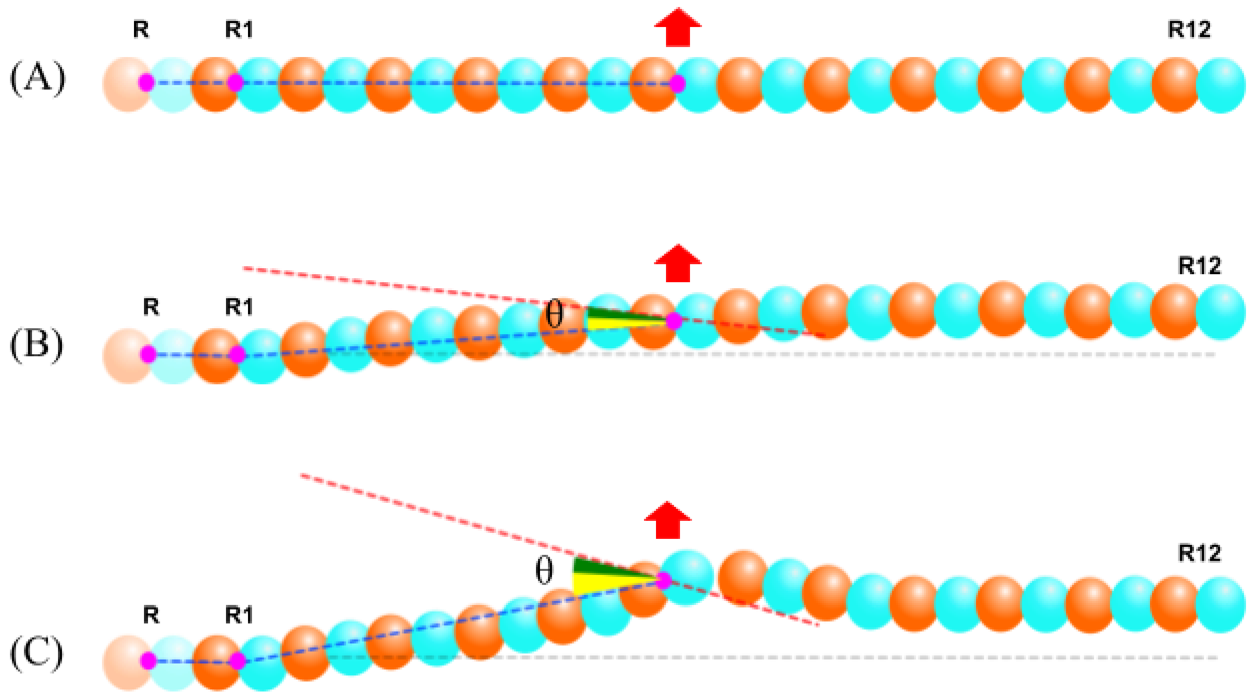


**Fig. S22:** Representation of bending angles measurements for a PF with free plus end. Orange and cyan spheres represent α and β tubulin monomers. R is the frozen dimer moved by 50 Å towards the minus end, R1 - fixed dimer and R12 - dimer on free plus end. The red arrow indicates the pulling point on PF and magenta indicates the center of mass of the respective dimers. (A) initial configuration (frame 0), (B) intermediate configuration as a result of pulling and (C) configuration corresponding to the first longitudinal break in the PF. The resulting angles are multiplied by 2 since the original angles do not cover the full MT.
